# Systemic inflammation accelerates neurodegeneration in a rat model of Parkinson’s disease overexpressing human alpha synuclein

**DOI:** 10.1101/2024.01.30.577912

**Authors:** Mariangela Massaro Cenere, Marta Tiberi, Emanuela Paldino, Sebastian Luca D’Addario, Mauro Federici, Cecilia Giacomet, Debora Cutuli, Alessandro Matteocci, Francesca Cossa, Beatrice Zarrilli, Nicolas Casadei, Ada Ledonne, Laura Petrosini, Nicola Berretta, Francesca Romana Fusco, Valerio Chiurchiù, Nicola B. Mercuri

**Affiliations:** Department of Experimental Neuroscience, Santa Lucia Foundation IRCCS, Rome, Italy; Department of Systems Medicine, University of Rome Tor Vergata, Rome, Italy; Aligning Science Across Parkinson’s (ASAP) Collaborative Research Network, Chevy Chase, MD, United States; Laboratory of Resolution of Neuroinflammation, Santa Lucia Foundation IRCCS, Rome, Italy; Laboratory of Neuroanatomy, Santa Lucia Foundation IRCCS, Rome, Italy; Department of Psychology, Sapienza University of Rome, Rome, Italy; PhD program in Immunology, Molecular Medicine and Applied biotechnologies, University of Rome Tor Vergata, 00133 Rome, Italy; Institute of Medical Genetics and Applied Genomics, University of Tübingen, Tübingen, Germany; Institute of Translational Pharmacology, National Research Council, Rome, Italy

## Abstract

Increasing efforts have been made to elucidate how genetic and environmental factors interact in Parkinson’s disease (PD). In the present study, we assessed the development of symptoms on a genetic PD rat model that overexpresses human α-synuclein (*Snca*^+/+^) at a presymptomatic age, exposed to a pro-inflammatory insult by intraperitoneal injection of lipopolysaccharide (LPS), using immunohistology, high-dimensional flow cytometry, constant potential amperometry, and behavioral analyses. A single injection of LPS into WT and *Snca^+/+^* rats triggered long-lasting increase in the activation of pro-inflammatory microglial markers, monocytes, and T lymphocytes. However, only LPS *Snca*^+/+^ rats showed dopaminergic neuronal loss in the *substantia nigra pars compacta* (SNpc), associated with a reduction in the release of evoked dopamine in the striatum. No significant changes were observed in the behavioral domain.

We propose our double-hit animal as a reliable model to investigate the mechanisms whereby α-synuclein and inflammation interact to promote neurodegeneration in PD.

## INTRODUCTION

Parkinson’s disease (PD) is a progressive neurodegenerative pathology that affects more than 7 million people worldwide ^1^. It is characterized by motor symptoms (such as bradykinesia, rigidity, resting tremor and postural instability) and nonmotor symptoms (lack of motivation, anxiety, dementia, depression, hallucinations, sleep disorders, pain, and gastrointestinal dysfunction) ^2–4^. The main pathological hallmarks of PD are a slow and progressive loss of dopaminergic (DAergic) neurons in the *substantia nigra pars compacta* (SNpc), causing decreased synaptic outflow of DA in the striatum, and accumulation of misfolded neuronal α-synuclein inclusions, known as Lewy bodies (LBs) and Lewy neurites (LNs) ^4–6^.

Most cases of PD are sporadic. However, genome-wide association studies (GWAS) have identified more than 90 genetic risk loci ^7^, responsible for an overall heritable component of the disease estimated at around 35% ^8^. Missense mutations and overexpression of the α-synuclein gene (SNCA), one of the most common mutations in monogenic PD, are widely used to model PD in rodents; however, no single animal model replicates all pathogenic and clinical characteristics of PD and some of them fail to develop nigral DAergic neurodegeneration. Therefore, there is common agreement that environmental factors significantly contribute to the pathogenesis of PD. In fact, it is worth noting that patients exhibiting the same genetic mutation may not have similar clinical presentations, suggesting that there is a complex interaction between environmental, age-associated, and genetic factors responsible for the etiology of the disease ^9,10^.

It has been proposed that endotoxins may participate in the pathogenesis of PD ^11^. Lipopolysaccharide (LPS) is the most used endotoxin to elicit an acute immune response. It is a major component of the outer membrane of Gram-negative bacteria and a potent inducer of inflammation in peripheral tissues and the central nervous system through the activation of toll-like receptor 4 (TLR4). Multiple studies support LPS treatment as an experimental model to reproduce PD-like symptoms in the animal. Progressive degeneration of DAergic neurons and microglial cell activation are reported after intraperitoneal injection (ip) of LPS in *wild-type* animals ^12–15^. Repeated injections of LPS over four consecutive days in 3-month-old male mice triggered a reduction in the number of neurons positive for SNpc tyrosine hydroxylase (TH +) and NeuN-positive neurons 19 days after the first injection of LPS ^15^; however, no further cellular reduction was found 36 days post-injection ^16^. The change in cytokine production, with increased anti-inflammatory and reduced pro-inflammatory cytokines, indicates that at later stages, animals develop cellular and molecular strategies to stop the ongoing neurodegenerative process ^16^. In line with this, TH protein levels decreased in the SN of Wistar rats 15 days after a single dose of LPS, followed by recovery 30 and 60 days post-injection ^17^.

In the present investigation, we tested the hypothesis that the development of PD occurs through a double-hit mechanism involving the combined interaction of elevated endotoxin and aggregated α-synuclein, with the result of neuronal degeneration in SNpc. To this end, we used rats overexpressing human α-synuclein (*Snca^+/+^*) characterized by a prodromal asymptomatic phase, with the first functional signs of DAergic pathology starting at four months of age, later resulting in a 25-27% depletion of TH and soma-dendritic neuronal modification at 12 months, accompanied by altered firing properties of DAergic neurons ^18–21^. In this animal model, we induced systemic inflammation at a presymptomatic age (2 months) through a single intraperitoneal injection of LPS, and we evaluated the effects on neuroinflammation and the DAergic system three months later.

## RESULTS

### Time course of sickness behavior and body weight

It should be noted that the doses of LPS usually used to induce a sickness response in mice or rats are about 10,000-100000 times higher than those administered to humans. This could be based on the inherent differences in the immune systems of laboratory rodents and humans. Consequently, higher doses of LPS are often necessary to induce an immune response in rodents ^22–24^. Furthermore, different LPS administration and dosage routes have been used to model specific characteristics of PD or other neurodegenerative diseases ^25,26^. Here, a dose of 5mg/kg of LPS was administered, as it is known that a single systemic administration of this dosage via the intraperitoneal cavity in wild-type male mice can induce chronic neuroinflammation and delayed progressive degeneration of TH^+^ neurons in the SN with onset around 7 months post-treatment^12–14,27–29^. To investigate whether a combinatorial effect of α-synuclein and LPS could lead to accelerated neurodegeneration, we looked at an earlier age.

Since LPS stimulates the release of several pro-inflammatory cytokines that are associated with fever, sickness behavior, and body weight loss ^30,31^, we first measured sickness signs and body weight over time (2h, 18h, 24h, 42h, 48h, 72h, 96h, and 168h) after injection of LPS ip (Supplementary Fig.1a-d). No signs of sickness were observed in saline (SAL)-treated WT and *Snca*^+/+^ rats treated with saline (SAL), except for slightly decreased exploration and locomotor activity in 10% of *Snca*^+/+^ rats (score =2) 24h after injection of saline. The LPS-treated groups exhibited clear signs of illness compared to controls. The maximum score was reached at 24h, when the rats manifested decreased locomotion and exploratory behavior, curled body posture, closed eyes, piloerection, and irregular fur (Supplementary Fig.1a, b). SAL *Snca*^+/+^ rats increased their body weight by 20 g ± 6 g over seven days, while LPS injection resulted in acute weight loss. Post hoc analysis demonstrated that LPS *Snca*^+/+^ rats lost weight in the first 72 h compared to SAL-treated rats, decreasing their body weight by 42 g ± 5 g at 18 h and then gradually recovering weight during the following weeks (Supplementary Figure 1c, d).

### Effects of peripheral LPS on long-lasting neuroinflammation

We next sought to investigate whether a single systemic dose of LPS injection was sufficient to cause long-lasting inflammation in both SNpc and the striatum, using the sophisticated approach of high dimensional flow cytometry, allowing us to identify all the different brain resident immune cells and peripheral immune cells (Fig.1a and Supplementary Fig. 2a-b).

**Figure 1.**
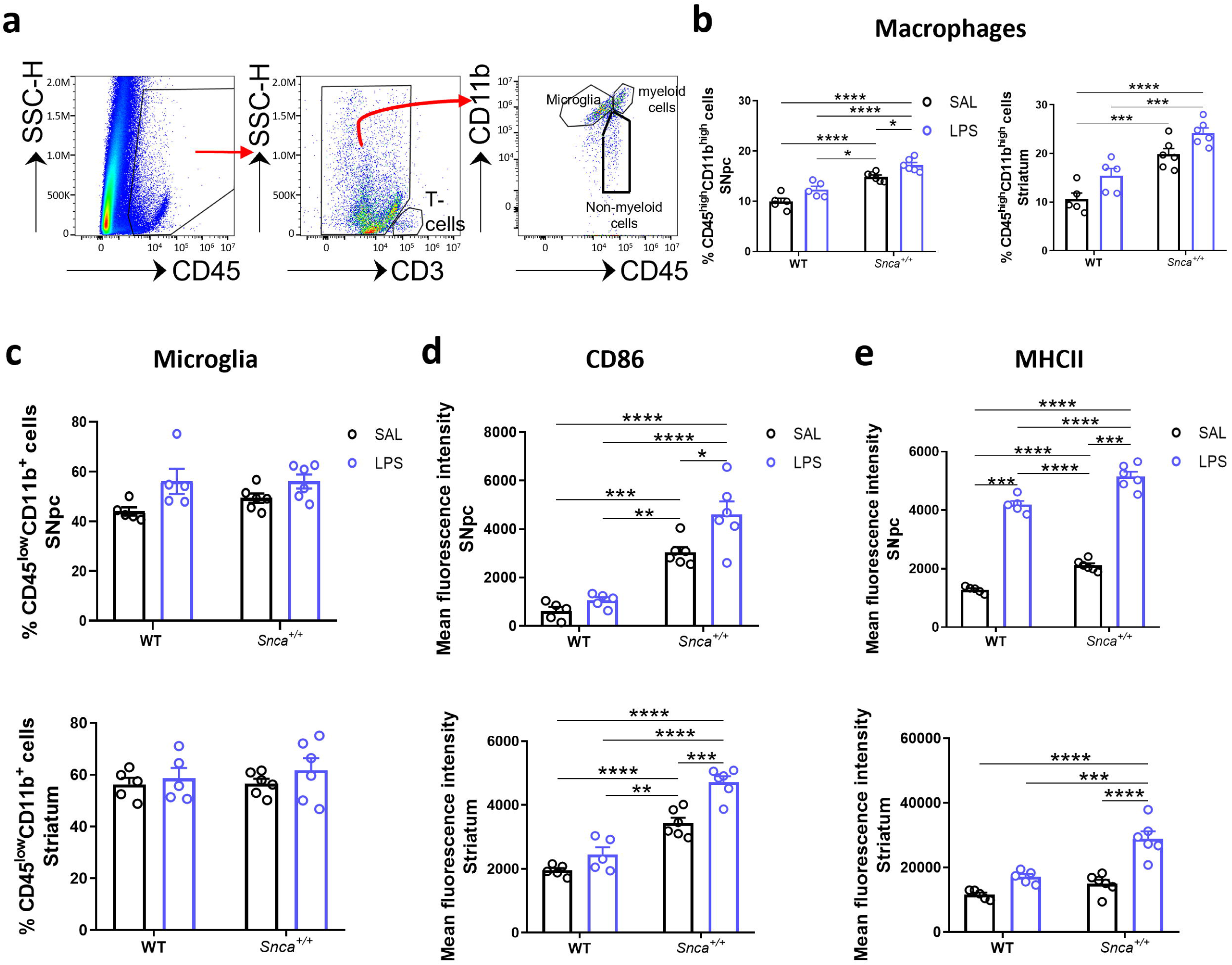
Monocyte and microglial activation in CNS 3 months after LPS administration. **a)** Representative flow cytometry dot plots of isolated cells illustrating the gating strategy to determine profiles based on CD45^low^CD11b^+^ for microglia, CD45^high^CD11b^high^ macrophages for the central nervous system (CNS)-associated phagocytes, and CD45^high^CD11b− cells. **b)** Percentage of CD45^high^CD11b^high^ cells in the SNpc (left) and in the striatum (right). Error bars represent ±SEM. (n=5 SAL WT, n=5 LPS WT, n=6 SAL *Snca^+/+^*, n=6 LPS *Snca^+/+^*; Ordinary two-way ANOVA for Genotype vs Treatment; SNpc: Interaction, F (1, 18) = 0.003483; Genotype, F (1, 18) = 79.06, p<0.001; Treatment, F (1, 18) = 18.48, p= 0.0004; *p<0.05, ****p<0.0001, with Bonferroni’s *post hoc* multiple comparisons test; striatum: Interaction, F (1, 18) = 0.02436; Genotype, F (1, 18) = 52.37, p<0.0001; Treatment, F (1, 18) = 13.76, p=0.0016; ***p<0.001, ****p<0.0001, with Bonferroni’s *post hoc* multiple comparisons test). **c)** Percentage of CD45^low^CD11b^+^ microglia in the SNpc (top) and in the striatum (bottom). Error bars represent ±SEM. (n=5 SAL WT, n=5 LPS WT, n=6 SAL *Snca^+/+^*, n=6 LPS *Snca^+/+^*; Ordinary two-way ANOVA for Genotype vs Treatment; SNpc: Interaction, F (1, 18) = 0.7841; Genotype, F (1, 18) = 0.7550; Treatment, F (1, 18) = 9.796, p=0.0058; striatum: Interaction, F (1, 18) = 0.1473). **d)** Mean fluorescence of CD86 both in the SNpc (top) and striatum (bottom). (n=5 SAL WT, n=5 LPS WT, n=6 SAL *Snca^+/+^*, n=6 LPS *Snca^+/+^*; Ordinary two-way ANOVA for Genotype vs Treatment; SNpc: Interaction, F (1, 18) = 2.858; Genotype, F (1, 18) = 79.88, p<0.0001; Treatment, F (1, 18) = 9.063, p=0.0075; *p<0.05, **p<0.01, ***p<0.001, ****p<0.0001, with Bonferroni’s *post hoc* multiple comparisons test; striatum: Interaction, F (1, 18) = 4.848, p= 0.0410; Genotype, F (1, 18) = 111.9, p<0.0001; Treatment, F (1, 18) = 25.47, p<0.0001; **p<0.01, ***p<0.001, ****p<0.0001, with Bonferroni’s *post hoc* multiple comparisons test). **e)** Mean fluorescence of MHC-II both in the SNpc (top) and striatum (bottom). (n=5 SAL WT, n=5 LPS WT, n=6 SAL *Snca^+/+^*, n=6 LPS *Snca^+/+^*; Ordinary two-way ANOVA for Genotype vs Treatment; SNpc: Interaction, F (1, 18) = 0.2921; Genotype, F (1, 18) = 56.27, p<0.0001; Treatment, F (1, 18) = 623.2, p<0.0001; ***p<0.001, ****p<0.0001, with Bonferroni’s *post hoc* multiple comparisons test; striatum: Interaction, F (1, 18) = 7.674, p=0.0126; Genotype, F (1, 18) = 24.36, p= 0.0001; Treatment, F (1, 18) = 40.04, p<0.0001; ***p<0.001, ****p<0.0001, with Bonferroni’s *post hoc* multiple comparisons test).

We observed that the percentage of CD45^high^CD11b^high^ monocytes/macrophages was higher in *Snca^+/+^* rats compared to WT rats, both in the SNpc and in the striatum. This percentage was even higher in the LPS-treated groups, especially *Snca^+/+^* rats (Fig.1b).

On the other hand, no significant changes were observed in the percentage of the resident myeloid cells of the brain, *i.e.,* CD45^low^CD11b^+^ microglia in the LPS-treated groups compared to the SAL-treated group (Fig.1c). Similar results were obtained with unbiased stereological cell count; no significant increase in the number of IBA1^+^cells was found in SNpc (Supplementary Fig.3a-b and Supplementary Table 1) and the striatum of LPS *Snca^+/+^* compared to SAL *Snca^+/+^* rats (Supplementary Fig.3e-f and Supplementary Table2).

Although the microglial cell count did not change, next we sought to further characterize the activation state of CD45^low^CD11b^+^ microglia by evaluating the expression of pro-inflammatory M1-like CD86 and MHC-II (Fig.1d). In WT rats, CD86 was not significantly altered in the two isolated brain regions three months after LPS administration (Fig.1d). MHC-II was significantly increased in the LPS WT group compared to the SAL-treated groups in SNpc (Fig.1e, top) but not in the striatum (Fig.1e, bottom). On the other hand, CD86 expression levels were higher in *Snca^+/+^* rats in both SNpc and striatum regardless of treatment and were even higher after LPS stimulation (Fig.1d). In contrast, MHC-II levels were increased in the *Snca^+/+^* rats compared to WT rats only in the SNpc (Fig.1e, top) and such levels were even higher after LPS treatment in both brain areas (Fig.1e).

Since the resting / activation state is associated with its morphological changes, we next performed a Sholl analysis of IBA1^+^ cells. In particular, even if no differences in IBA1^+^ microglia cells were observed, they showed decreased complexity, shorter branching and decreased number of intersections, endings, and nodes in LPS *Snca^+/+^*compared to the SAL-treated group in SNpc (Supplementary Figure 3c-d). In the striatum, only slight differences were observed in the number of intersections, and dendritic length (Supplementary Fig. 3gh)

These data suggest that a single systemic dose of LPS triggers a neuroinflammatory response in the SNpc and striatum that likely involves microglial activation.

### Effects of peripheral LPS on leukocyte levels

Given that peripheral immune cells are also present in brain tissue and contribute to disease pathology ^32^, we investigated whether a single systemic administration of LPS affects peripheral immune cells in the SNpc and the striatum. To this end, the rest of the CD45^high^CD11b^low^ cells (Fig.1a) were further gated by high dimensional flow cytometry to identify CD3^+^ T-lymphocytes, CD45RA^+^ B-lymphocytes and CD161^+^ NK cells, according to their signature identifying markers. We observed that only T-lymphocytes were increased in the SNpc of LPS *Snca^+/+^* rats compared to SAL- and LPS-treated WT rats in terms of both cell percentages (Fig.2a, top) and cell count (Supplementary Fig.2c). To check whether T cells were present in SNpc and striatal parenchyma, immunostainings of coronal brain sections of these areas were performed and few CD3^+^ cells were observed near TH^+^ neurons in *Snca^+/+^* rats SNpc (Supplementary Figure 4a) and in LPS-treated rats striatal sections (Supplementary Figure 4b). CD3 + cells were observed in WT rats, nor in the striatum of SAL-treated animals. These data qualitatively suggest the presence of T-cells in some brain regions, despite the impossibility to discriminate between infiltrating immune cells and those that might randomly remain in blood vessels during tissue collection (see Methods). It is worth noting that we measured very few T-cells by flow cytometry, with the mean number of T-cells estimated to be between 10 and 120 in WT SAL animals (SNpc or striatum), between 40 and 180 in WT LPS animals (SNpc or striatum), between 60 and 160 in Snca^+/+^ SAL animals (SNpc or striatum) and between 300 and 580 in SNCA^+/+^-LPS animals (SNpc or striatum) (Supplementary Fig.2c). These numbers reflect the presence of T-cells in the entire dissociated tissue of the dissected areas, while immunofluorescence was performed in 3 slices for each region.

**Figure 2.**
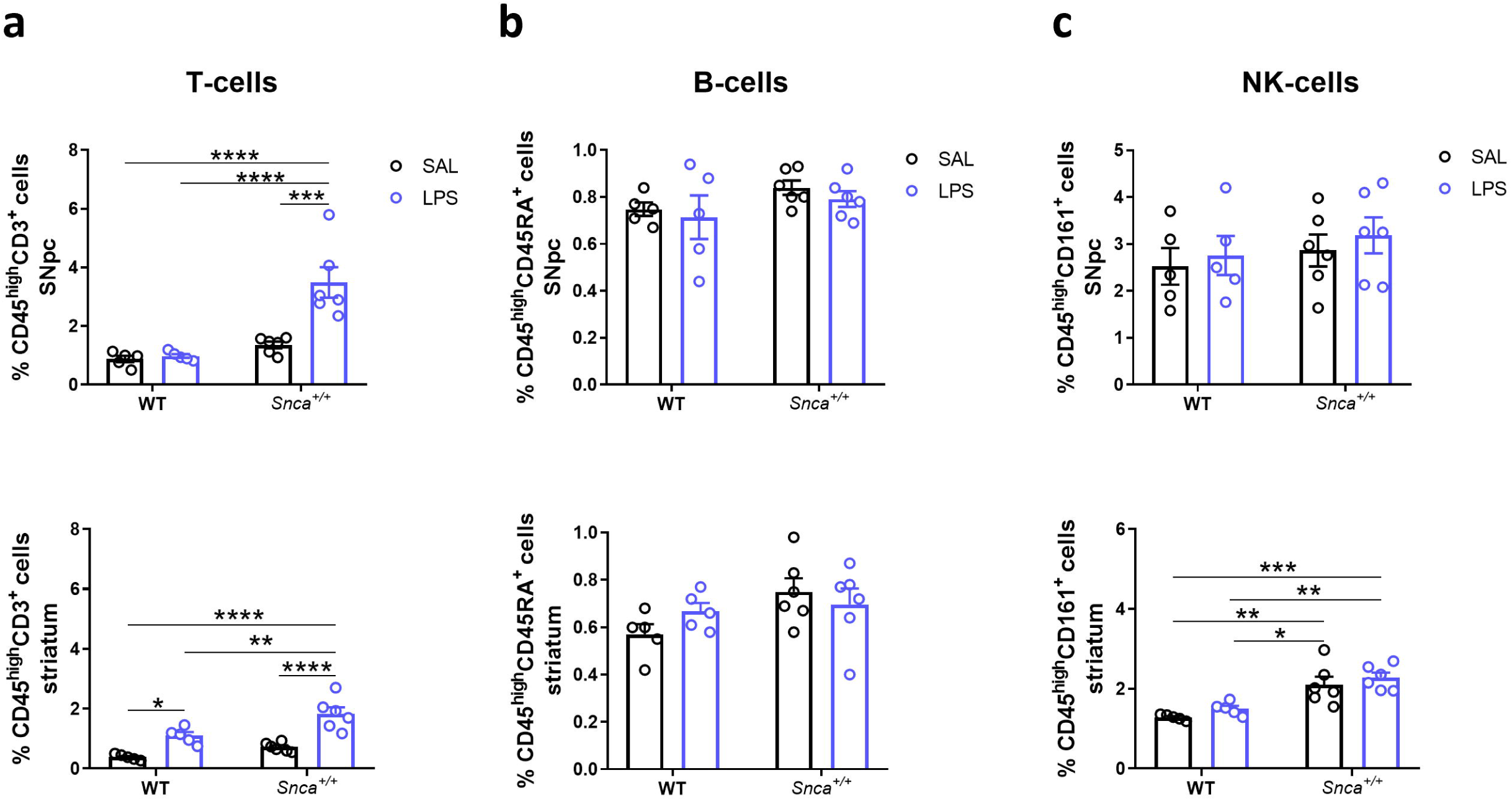
Peripheral immune cells into the SN and striatum 3 months after LPS administration. **a)** Bar plots of the percentage of CD45^high^CD3^+^ T cells in the SNpc (top) and striatum (bottom). (n=5 SAL WT, n=5 LPS WT, n=6 SAL *Snca^+/+^*, n=6 LPS *Snca^+/+^*; Ordinary two-way ANOVA for Genotype vs Treatment; SNpc: Interaction, F (1, 18) = 11.58, p=0.0032; Genotype, F (1, 18) = 25.14, p<0.0001; Treatment, F (1, 18) = 14.07, p=0.0015; ***p<0.001, ****p<0.0001, with Bonferroni’s *post hoc* multiple comparisons test; striatum: Interaction, F (1, 18) = 2.041; Genotype, F (1, 18) = 15.05, p= 0.0011; Treatment, F (1, 18) = 44.63, p<0.0001; *p<0.05, **p<0.01, ****p<0.0001, with Bonferroni’s *post hoc* multiple comparisons test). **b)** Bar plots of the percentage of CD45^high^CD45RA^+^ B cells in the SNpc (top) and striatum (bottom). (n=5 SAL WT, n=5 LPS WT, n=6 SAL *Snca^+/+^*, n=6 LPS *Snca^+/+^*; Ordinary two-way ANOVA for Genotype vs Treatment; SNpc: Interaction, F (1, 18) = 0.02012; striatum: Interaction, F (1, 18) = 1.916). **c)** Bar plots of the percentage of CD45^high^CD161^+^NK cells in the SNpc (top), while, in the striatum (bottom), a significant genotype effect in *Snca^+/+^* compared to WT groups is shown, independent from treatment. (n=5 SAL WT, n=5 LPS WT, n=6 SAL *Snca^+/+^*, n=6 LPS *Snca^+/+^*; Ordinary wo-way ANOVA for Genotype vs Treatment; SNpc: Interaction F (1, 18) = 0.01421; striatum: Interaction, F (1, 18) = 0.01596; Genotype, F (1, 18) = 33.17, p<0.0001; Treatment, F (1, 18) = 2.033; *p<0.05, **p<0.01, ***p<0.001, with Bonferroni’s *post hoc* multiple comparisons test).

No changes were found in the percentage of B-lymphocytes and NK cells (Fig.2b-c). On the other hand, in the striatum, increased percentages of T lymphocytes were also observed in WT rats after LPS treatment (Fig.2a, bottom), along with a statistically significant increase in NK cells in *Snca^+/+^* rats compared to WT rats, but not following LPS treatment (Fig.2c, bottom panel). Since we reported an increase in T lymphocytes within the brain of *Snca^+/+^* rats and recent evidence supports a key role for T cells in the pathogenesis of PD ^33^, we further investigated the immunophenotype of peripheral blood mononuclear cells (PBMC) (Supplementary Fig.5a) to assess whether differences in immune cells into brain tissue reflect changes in peripheral blood. Although no significant changes were observed in the percentage of B cells, NK cells, and granulocytes (respectively, Supplementary Fig.5b-d), monocytes were higher only in LPS *Snca*^+/+^ rats compared to the other experimental groups (Supplementary Fig.5e), while T cells were significantly increased in both genotypes after LPS stimulation (Supplementary Fig.5f).

### Effects of peripheral LPS on alpha-synuclein accumulation

To understand how LPS-induced microglial activation might influence α-synuclein accumulation in our Parkinson’s disease model, we assessed phosphorylated α-synuclein (pSyn S129) in various brain regions of WT and *Snca^+/+^*rats three months after SAL or LPS injection. In *Snca^+/+^*rats, α-synuclein is overexpressed in several areas of the brain with the accumulation of different pathological forms over time ^18–20^. We observed widespread immunopositive signals for pSyn (S129) in various brain regions (Fig.3a, d). In particular, our findings revealed a sublet increase in pSyn S129 immunoreactivity within SNpc of LPS *Snca^+/+^* rats compared to those injected with saline (Fig.3a). In particular, this effect was not observed in WT animals, suggesting a possible interaction between LPS exposure and the pre-existing α-synuclein pathology in *Snca^+/+^*rats. pSyn (S129) levels were not altered in the Ventral tegmental area (VTA), another DAergic nucleus adjacent to SNpc, in response to LPS administration (Fig.3b). This suggests regional specificity in the LPS-mediated effects on α-synuclein phosphorylation levels. In the striatum, while both SAL- and LPS-treated *Snca^+/+^* rats displayed higher pSyN (S129) levels compared to WT rats, a difference emerged between SAL- and LPS-treated *Snca^+/+^* rats, with higher levels in SAL *Snca^+/+^* than in LPS *Snca^+/+^* rats (Fig.3c).

**Figure 3.**
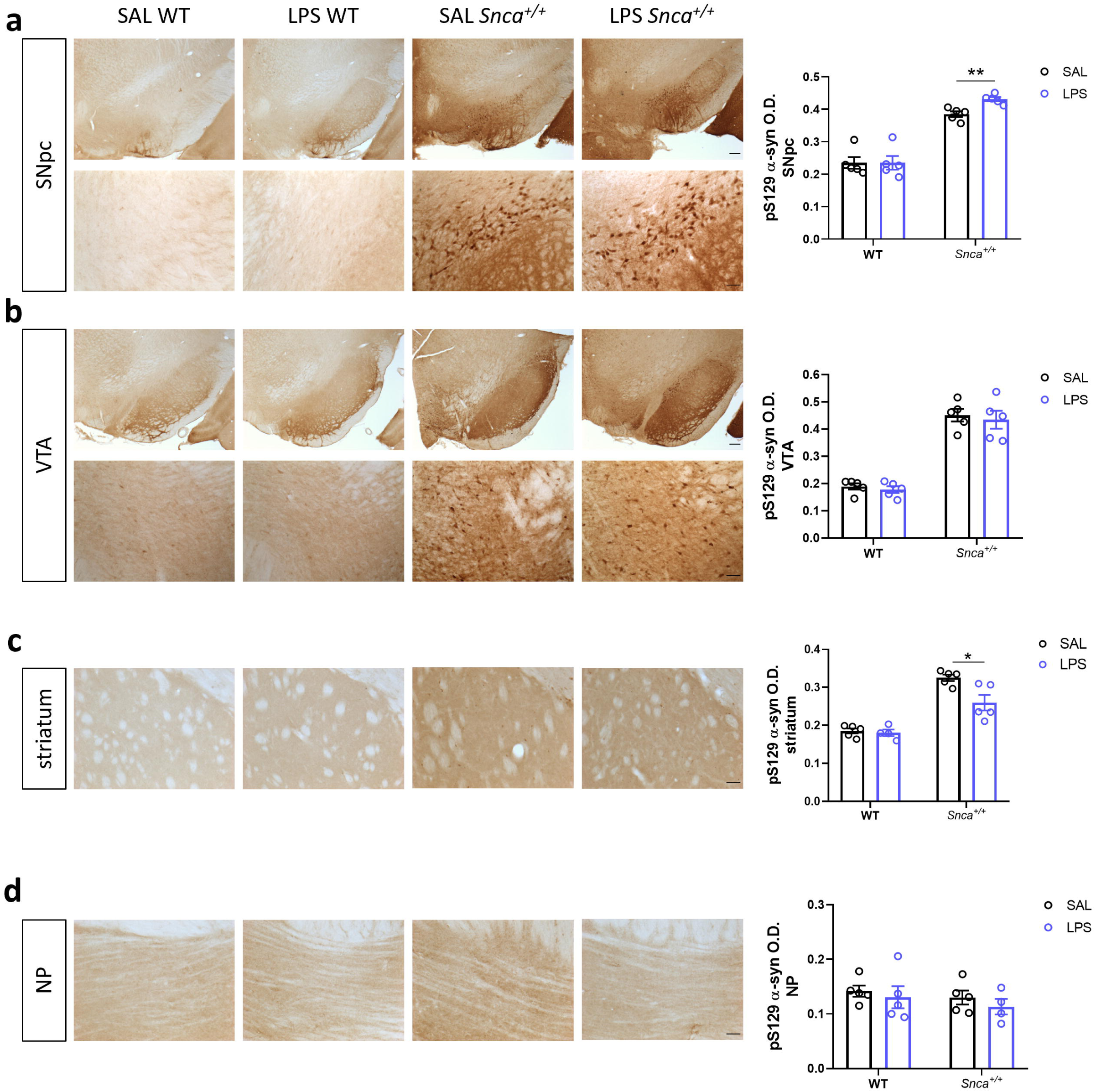
LPS accelerates alpha synuclein accumulation in the SN of *Snca^+/+^* rats. **a)** Left, representative immunostaining of pSyn (S129) in the SNpc (top, scale: 200µm) and related micrograph (bottom, scale: 50μm). Right, densitometric quantification of pSyn (S129) in SNpc. Each point represents mean O.D. values per animal ±S.E.M. (n=5 rats/group; Kruskal-Wallis test: p= 0.0011 (approximate), **p= 0.0079, with Mann Whitney test). **b)** Left, representative immunostaining of pSyn (S129) in the VTA (top, scale: 200µm) and micrograph (bottom, scale: 50μm). Right, densitometric quantification of pSyn (S129) in VTA. Each point represents mean O.D. values per animal ±S.E.M. (n=5 rats/group; Ordinary Two-way ANOVA for Genotype vs Treatment: Interaction, F (1, 16) = 0.01289; Genotype, F (1, 16) = 138.5, ****p<0.0001; Treatment, F (1, 16) = 0.3907). **c)** Left, representative immunostaining of pSyn (S129) in the striatum (scale: 50μm) and densitometric quantification, right. Each point represents mean O.D. values per animal ±S.E.M. (n=5 rats/group; Ordinary Two-way ANOVA for Genotype vs Treatment: Interaction, F (1, 15) = 5.632, p= 0.0314; Genotype, F (1, 15) = 73.16, p<0.0001; Treatment, F (1, 15) = 7.504, p= 0.0152; *p= 0.122, with Bonferroni’s post hoc multiple comparisons test). **d)** Left, representative immunostaining of pSyn (S129) in the NP (scale: 50μm) and densitometric quantification, right. Each point represents mean O.D. values per animal ±S.E.M. (n=5 rats/group; Ordinary Two-way ANOVA for Genotype vs Treatment: Interaction, F (1, 15) = 0.03596)

Finally, the pontine nuclei, a brain region known to be unaffected by α-synuclein accumulation in this model^19^, served as a negative control. As expected, the immunoreactivity of pSyn (S129) remained consistent across all groups (Fig.3d).

These findings suggest that peripheral exposure to LPS may influence the phosphorylation and potential aggregation of α-synuclein, particularly within the SNpc of the *Snca^+/+^* model. This effect appears to be specific to brain regions already prone to α-synuclein pathology, highlighting a potential role for neuroinflammation in disease progression.

### Effects of peripheral LPS on nigrostriatal alterations in Snca^+/+^rats

To test the hypothesis that endotoxin may enhance nigral DAergic neuron degeneration in *Snca*^+/+^ rats, we evaluated immunohistological changes in nigral neurons three months after injection via the unbiased stereological cell count, staining midbrain coronal sections with TH as a DAergic marker. The unbiased stereological count confirmed the absence of TH^+^ neuronal loss in SAL *Snca^+/+^*compared to SAL WT rats (Fig.4a-b), as previously reported ^19^. Three months after a single LPS i.p. injection, both WT and *Snca^+/+^*groups showed a statistically significant decreased number of TH^+^ neurons in the SNpc, by almost 28% (±7% SEM) and 49% (±8% SEM) compared to SAL WT rats (Fig.4b). Stereological cell counts did not show any differences in the number in the VTA, the adjacent DAergic nucleus that remains spared during the initial stages of progression of PD (Fig. 4c). This supports the preferential vulnerability of nigral DAergic neurons under prolonged α-syn overexpression, in line with evidence in the same aged rat model ^19^.

**Figure 4.**
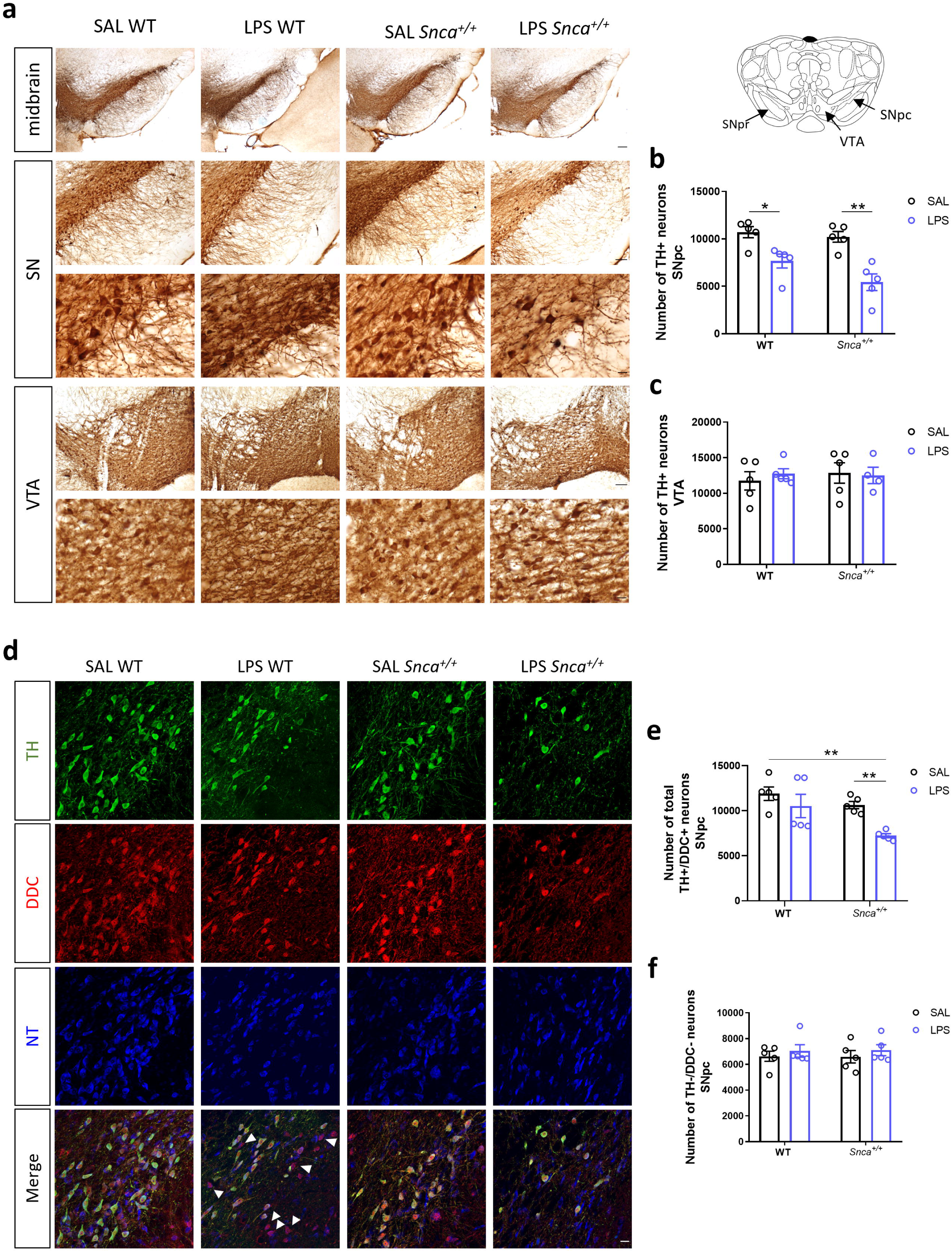
Morphological alterations in SNpc induced by LPS. **a)** Representative coronal images midbrain, SN, and VTA slices from 3,3’-diaminobenzidine (DAB)-stained TH^+^ neurons in WT and *Snca^+/+^*rats treated with saline (SAL) or Lipopolysaccharide (LPS) at 3 months after injection (scale: 100µm). **b)** Unbiased stereological TH^+^ cell count plot in the SNpc. Each point represents actual values ±SEM. (n=5 rats/group; Kruskal-Wallis test: p= 0.0031 (approximate); *p= 0,0159, **p= 0,0079, with Mann Whitney test). **c)** Unbiased stereological TH^+^ cell count plot in the VTA. Each point represents actual values ±SEM. (n=5 rats/group; (n=5 rats/group; Ordinary two-way ANOVA for Genotype vs Treatment: Interaction, F (1, 15) = 0.3154). **d)** Representative confocal images of stained SNpc sections for TH (green) and DDC (red) dopaminergic markers and NeuroTrace (NT, blue) in WT and *Snca^+/+^* rats treated with saline (SAL) or Lipopolysaccharide (LPS) at 3 months after injection (scale: 20µm). White arrowheads show TH^-^/DDC^+^/NT^+^ neurons (magenta). **e)** Unbiased stereological count of total TH^+^/DDC^+^ neurons in SNpc. Each point represents actual values ±SEM. (n=5 rats/group; Kruskal-Wallis test: p=0.0082 (approximate); **p<0.01, with Mann Whitney test). **f)** Unbiased stereological count of total TH^-^/DDC^-^ neurons in SNpc. Each point represents actual values ±SEM (n=5 rats/group; Kruskal-Wallis test: p=0.9016 (approximate)).

Since a study by Heo and colleagues ^34^ demonstrated that some SNpc DAergic neurons could lose their TH expression and become so-called dormant neurons, we tested whether the observed decrease in TH^+^ neurons in LPS-treated rats could reflect down-regulation of TH rather than an actual neuronal loss, using DOPA decarboxylase (DDC) as an additional DAergic marker. We found TH-negative neurons (TH-) that were DDC-positive (DDC^+^) mainly in LPS WT, but not in LPS *Snca*^+/+^ rats, and the statistically significant difference of TH^+^ neurons in LPS WT rats was lost when comparing the total number of dopaminergic neurons (sum of TH^+^/DDC^+^ and TH^-^/DDC^+^) (Fig.4d-e).

Furthermore, no statistically significant differences were found in the neuronal cell count among the four experimental groups(Fig.4f).

Collectively, these data support the hypothesis of a TH downregulation in LPS WT rats, meaning that some SNpc DAergic neurons lose their TH-phenotype and could be surviving or bound to die. In contrast, a real and selective loss of DAergic neurons occurs only in LPS *Snca*^+/+^ rats.

SNpc DA neurons have extensive dendritic arborization that extends ventrally towards the SN pars reticulata (SNpr). In *Snca^+/+^* rats injected with LPS, this TH^+^ dendritic projection showed changes in morphology and density. Dendrites from the remaining DAergic neurons appeared truncated with a distorted morphology (swollen and bubble-shaped) compared to the other experimental groups, showing a smooth and lengthened profile (Fig.5g). Optical densitometry showed a significant loss of TH^+^ dendrites at three months after LPS injection, compared to SAL-treated groups (Fig.5h).

**Figure 5.**
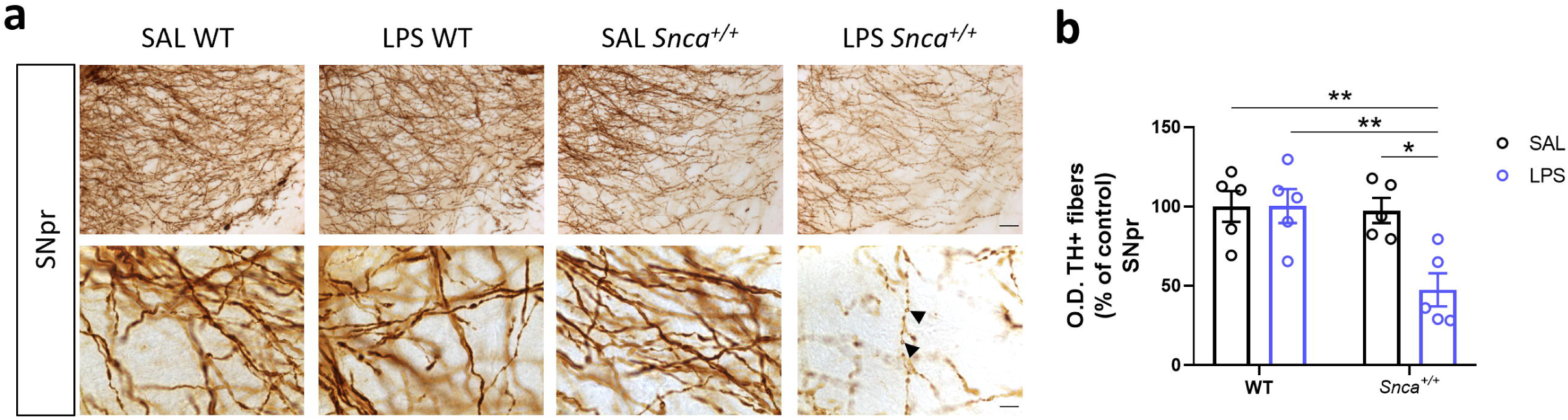
Morphological alterations in SNpr induced by LPS in *Snca^+/+^* rats. **a)** Representative image of coronal midbrain slice from DAB-stained SNpr TH^+^ fibers in WT and *Snca^+/+^* rats treated with SAL or LPS (scale: 50µm) and related micrographs (scale: 10µm). Black arrowheads show swollen TH^+^ dendrites morphology. **b)** SNpr TH^+^ fibers densitometric analysis. Values are expressed as means of each animal normalized to the total mean value of SAL WT group ± SEM. (n=5 rats/group; Ordinary two-way ANOVA for Genotype vs Treatment: Interaction, F (1, 16) = 6.614, p=0.0205; Genotype, F (1, 16) = 8.053, p=0.0119; Treatment, F (1, 16) = 6.497, p=0.0214; *p<0.05, **p<0.01, with Bonferroni’s *post hoc* multiple comparisons test).

Densitometric analysis of multiple sections of DAergic terminals in the dorsal striatum did not show a significant decrease in TH immunoreactivity in LPS-treated rats compared to SAL-treated groups, despite the decrease in SNpc TH^+^ neurons and altered dendritic arborization (Fig.6d). Furthermore, constant potential amperometry measurements of DA outflow in the dorsolateral striatum revealed a decrease in the evoked DA amplitude in LPS *Snca^+/+^*rats compared to SAL-treated rats (Fig.6a), with no change in half-decay time (192.7 ± 1.7 ms vs. 195.9 ± 1.9 ms, respectively). In previous studies in four-month-old naïve *Snca^+/+^*rats ^19^, we found a reduced DA outflow in the dorsal striatum in SAL *Snca*^+/+^ compared to WT rats. Here, we found a more significant decrease in DA release when combining genetic and endotoxic risk factors. No significant differences were found in LPS WT rats compared to SAL WT (Fig.6a) in either amplitude or half decay.

**Figure 6.**
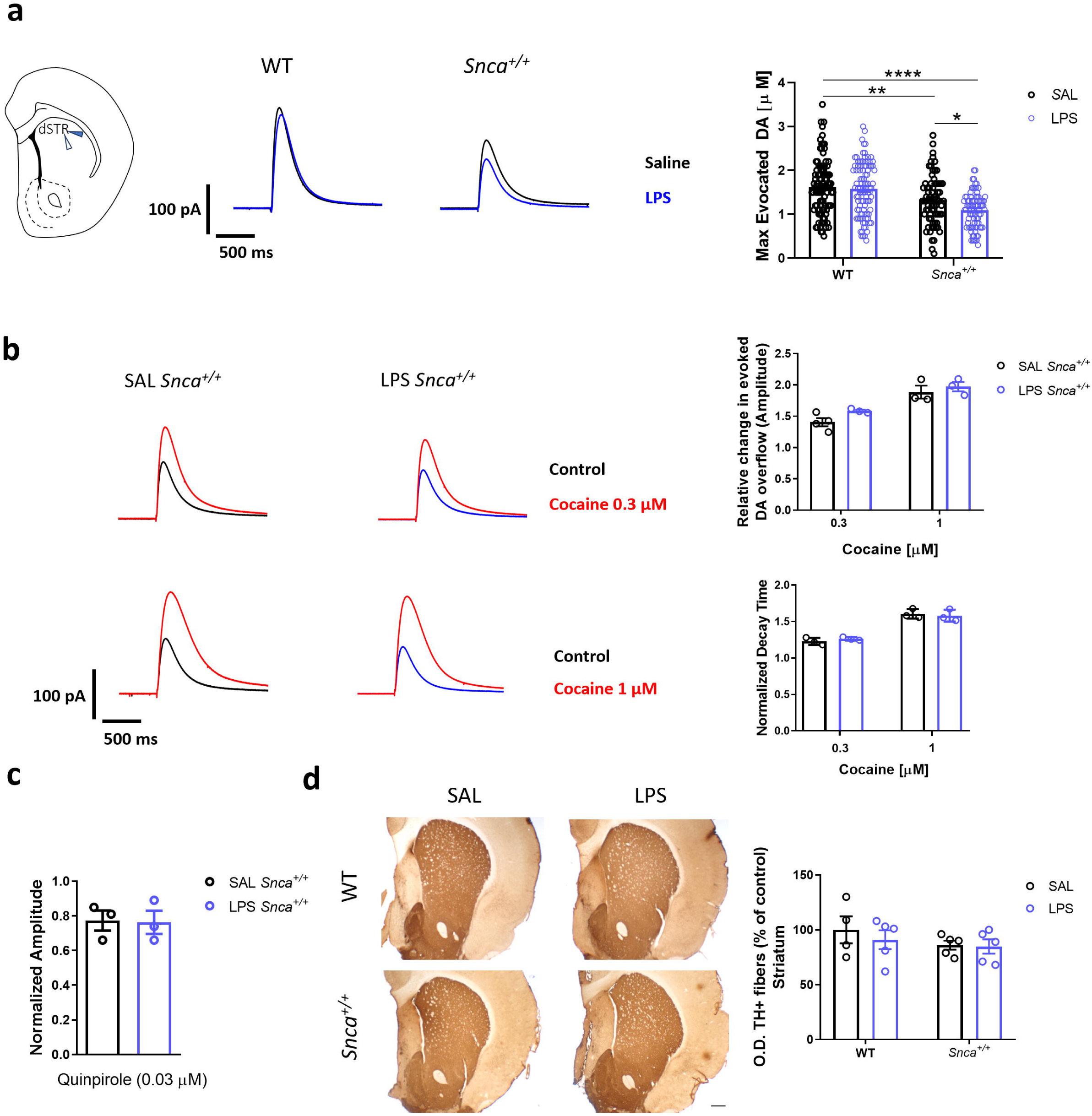
LPS induced functional and morphological alterations in the striatum. **a)** Left, **s**chematic representation showing the placement of the stimulating (blue arrowhead) and carbon fiber (white arrowhead) used for the recordings in the dorsal striatum. Middle, representative traces (scale: 100 pA; 500□ms) and evoked DA concentration in the dorsal striatum at 3 months after treatment in WT and *Snca^+/+^* rats recorded with constant potential amperometry. Right, bar plot showing max evoked DA outflow in *Snca^+/+^* rats. (Ordinary two-way ANOVA Genotype vs Treatment; Interaction, F (1, 343) = 2.395; Genotype, F (1, 343) = 41.03, p<0.0001; Treatment, F (1, 343) = 5.763, p=0.0169; *p<0.05, **p<0.01, ****p<0.0001, with Tukey’s *post hoc* multiple comparisons test). **b)** Left, representative traces showing the effects of cocaine at different concentrations (top: 0.3μM; Bottom: 1μM) in SAL (left) and LPS (right) *Snca^+/+^* rats. At the top right, the bar plot indicates the effect of cocaine on DA overflow (n= 4 SAL *Snca^+/+^* cocaine [0.3μM], n=3 LPS *Snca^+/+^*cocaine [0.3μM], n= 3 SAL *Snca^+/+^* cocaine [1μM], n=3 LPS *Snca^+/+^* cocaine [1μM]; Ordinary two-way ANOVA Treatment vs Cocaine concentrations; Interaction, F (1, 9) = 0.3939). On the right bottom, the bar plot indicates the effect of cocaine on the decay phase of the DAergic signal. Data are presented as mean ± SEM. (n= 4 SAL *Snca^+/+^* cocaine [0.3μM], n=3 LPS *Snca^+/+^*cocaine [0.3μM], n= 3 SAL *Snca^+/+^* cocaine [1μM], n=3 LPS *Snca^+/+^*cocaine [1μM]; Ordinary two-way ANOVA Treatment vs Cocaine concentrations; Interaction, F (1, 9) = 0.1171). **c)** The bar plot shows the change in striatal DA release upon 5 min superfusion of quinpirole. Data are presented as mean ± SEM. (n= 3 rats/group; *t-*test, t=0.1126). **d)** Representative images of TH immunoreactivity of striatal coronal sections in WT and *Snca^+/+^* rats treated with SAL or LPS 3 months after treatment (scale bar: 500 μm). The values are expressed as means of each animal normalized to the total mean value of SAL WT group ± SEM. (n=5 rats/group; Ordinary two-way ANOVA for Genotype vs Treatment; Interaction, F (1, 15) = 0.2409).

Next, to further explore possible alterations of DA release, we analyzed the modifications of the DA transporter (DAT) function in *Snca^+/+^* rats via superfusion of striatal coronal slices with the DAT blocker cocaine. As expected, cocaine (0.3-1 μM) increased the amplitude of the amperometric signal in a concentration-dependent manner and increased its half-decay time ^35^. However, we did not find any statistically significant differences between the two experimental groups either at low (0.3 µM) or high (1 µM) concentrations of cocaine (Fig.6b). Additionally, we next investigated the function of DA autoreceptors (D2R) on DAergic transmission ^36,37^. To this end, we superfused striatal slices with a low dose of the D2-agonist quinpirole (0.03 µM). As expected, quinpirole reduced evoked DA efflux by almost 30%; however, no difference was observed in quinpirole effect on the DA signal between SAL *Snca^+/+^* and LPS *Snca^+/+^* rats (Fig.6c).

Possible functional alterations in DAergic transmission were also evaluated by assessing the animal’s general locomotor activity and depressive-like behavior via the rotarod and open field tests and the sucrose preference test, respectively. No significant differences were observed in these behavioral tests in the animals three months after a single LPS dose (Supplementary Fig.6a-c). Since the two-way ANOVA revealed a significant treatment effect on sucrose solution intake, we then performed Tukey’s multiple comparison *post hoc* test; however, no differences were noted between the four experimental groups (Supplementary Figure 6c).

## DISCUSSION

There is now a wide consensus that PD is a multisystem disorder that involves genetic and environmental risk factors. However, understanding the complex mechanisms underlying gene and environment interactions presents a significant challenge. Our present data propose human α-synuclein expressing rats injected with LPS as a reliable model to investigate how these two factors cooperate in the genesis and progression of neurodegenerative processes in the DAergic system typical of PD, pointing to inflammation as a connective element of this harmful cooperation.

A prodromal symptomatic phase usually characterizes PD animal models based on SNCA gene mutation and overexpression. Previously, we described the human α-synuclein overexpressing rat model, used in this study, showing a presymptomatic phase preceding early pathological signs occurring at four months ^19^. At this age, rats exhibit a significant reduction in striatal DA release despite the integrity of DAergic terminals and in the absence of nigral neuronal loss. These early alterations are paralleled by neuroinflammation in the CNS, with microglial activation and increased levels of pro-inflammatory cytokines in the cerebrospinal fluid (INFγ), while in blood granulocytes, T, and B cells are not altered ^19^.

Here, we showed that *Snca^+/+^* rats exhibit robust, long-lasting central and peripheral inflammatory activation when combined with an early event of acutely induced inflammation accompanied by sublet increased levels of pSyn (S129) in the SNpc.

Interestingly, enhanced nigral IBA1 expression was also reported in a transgenic mouse model expressing mutated α-synuclein, i.p. injected with LPS at the early symptomatic age ^38^. This study shows a delayed chronic and progressive degeneration of nigral TH^+^ neurons, with a more prominent effect five months after LPS injection. Another recently published study by Thi Lai and co-workers^39^ reported a higher IBA1-positive microglial density in response to a coinjection of LPS and α-synuclein preformed fibrils (PFF) in mice compared to LPS or PFF only 14 days after injection, accompanied by worsening of synucleinopathy.

Previous studies suggested different roles for microglia in the α-synuclein propagation. In fact, in a resting state, it could phagocytize and degrade α-synuclein, while an overload of α-synuclein may activate microglia through the Toll-like receptor 2 (TLR2) and, in turn, contribute to α-synuclein pathology by releasing pro-inflammatory cytokines promoting α-synuclein aggregation, cell-to-cell transfer, and neuronal death ^40–43^.

Furthermore, in a recent study, microglia, virally transduced to express the human SNCA mutant gene, developed a robust reactive state with phagocytic exhaustion and a toxic environment, with excessive production of oxidative and pro-inflammatory molecules, leading to progressive degeneration of DA neurons without endogenous α-synuclein aggregation ^44^.

Notably, our data suggest a marked microglia activation, polarizing towards a more pro-inflammatory phenotype in LPS *Snca^+/+^*rats in SNpc and striatum, accompanied by increased levels of monocytes and T-lymphocytes. On the contrary, LPS WT rats showed mild differences from SAL-treated WT, without macrophage differences and only increased levels of microglia MHC-II pro-inflammatory activation markers in the SN and few T lymphocytes in the striatum.

The role of T cells in upregulating microglia MHC-II expression has been demonstrated in viral-induced expressing α-synuclein rats. This appears to be an important step in dopamine neuronal loss in SNpc in response to α-synuclein^45^. The contribution of MHC-II to PD pathology is inferred from studies demonstrating that MHC null mice are resistant to dopaminergic degeneration under conditions of α-synuclein overexpression^46^. Importantly, a set of peptides derived from α-syn have been found to act as antigenic epitopes to further drive the responses of CD4 + and CD8 + cells in PD patients^33^.

Therefore, activated microglia by α-synuclein may induce the selective recruitment of peripheral immune cells that could induce further activation of microglia through the expression of MHC-II. This inflammatory state can create a molecular feed-forward vicious cycle between microglia and peripheral immune cells, mediating neuronal degeneration.

The increased level of T lymphocytes reported in the present study is of particular interest since T cells are known to play a key role in both the CNS and the periphery, leading to a profound imbalance of the immune network in patients with PD ^47^. T lymphocytes were reported in postmortem brain samples from PD patients ^29,30^ and a mouse model of PD ^48^; additionally, knockout of CD4 + T cells in a PD mouse model resulted in markedly reduced DAergic neuronal death ^48^. These data strengthen the hypothesis that T cells are crucial for neurodegeneration during PD and support the contribution of autoimmune-based pathogenesis for this disease ^25,31,32^. It is noteworthy that further studies will be warranted to better dissect which specific T cell subset is the most involved and has the highest pathogenic potential.

Since inflammation triggers neurodegeneration, we hypothesized that LPS could also affect the viability of the DAergic neurons. Our findings show that LPS injection in *Snca^+/+^* rats significantly degenerates DAergic neurons. This was observed at an age (5 months) when these animals do not normally present signs of neurodegenerative processes in the SNpc. Although the observed reduction in TH ^+^ neurons in both WT and *Snca^+/+^* rats occurs independently from α-synuclein, the decrease in DDC^+^ neurons that only occurred in LPS *Snca^+/+^* rats suggests that DAergic neuron death only happens in response to a double hit of genetic human α-synuclein overexpression and endotoxin-induced inflammation. The decrease in TH ^+^ neurons in LPS WT rats could reflect the down-regulation of TH. Indeed, Heo et al. defined neurons without TH as dormant DAergic cells that survive or are bound to die at later stages ^34^. Future studies are needed to assess whether these neurons will die in LPS-injected WT rats in later phases. It is also worth mentioning that a significant portion of TH^-^/DCC^+^ neurons was demonstrated in the SNpc of various animal models and post-mortem PD ^34^. Similar results were reported on the *parkin^-/-^* mice, another PD model, after six months of repeated low-dose intraperitoneal LPS injections. Both *wild-type* and *parkin*^-/-^ mice show substantial TH^+^ neuronal loss, but only in LPS-treated *parkin^-/-^* mice there is a significant reduction of NeuN-positive neurons in the SNpc ^44^.

Another important feature highlighted in the present study is the observed dystrophic TH^+^ dendritic arborizations, branching from SNpc surviving DAergic neurons towards the SNpr in LPS *Snca^+/+^* rats, anticipating by almost seven months similar alterations in the naϊve *Snca^+/+^* animals ^21^. Loss of neuronal complexity and the decreased dendritic arborization have been linked to α-synuclein overexpression in virally infected DAergic neurons *in vitro* and *in vivo*, before their eventual death ^49,50^ and similar to postmortem alterations in patients with PD ^51^.

The loss of DAergic neurons is associated with a greater extent of reduced evoked striatal DA release in LPS *Snca^+/+^* compared to SAL *Snca^+/+^* rats. However, no significant differences were observed in TH immunoreactivity in the striatum. In other genetic α-synuclein-based models, TH immunoreactivity in the striatum is unchanged in the early phases ^20,19,52^ and in the *parkin / -* PD model with repeated LPS administrations ^53^. A possible explanation for the lack of reduction in striatal TH immunoreaction despite DAergic neuronal loss in the SNpc could be the presence of compensatory upregulation of TH due to the sprouting of the surviving DAergic terminals. In addition to T lymphocytes, our data suggest an increase in NK cells in the striatum and this could explain its neuroprotective role in PD that has been proposed based on *in vitro* and *in vivo data suggesting* that it could clear α-synuclein aggregates ^54–58^, thus protecting the surviving DAergic terminals, at least in an initial phase of pathology progression.

The increased expression of activation markers in microglia like MHC-II and CD86, which provide, respectively, the first and costimulatory signals necessary for T cell activation, coupled with the increased percentages of peripheral T cells in both brain areas suggest the establishment of an immunological synapse in key areas involved in PD pathology which could drive and sustain neuroinflammation and lead to neurodegeneration.

According to our results, the decrease in DAergic neurons in the SNpc and evoked striatal DA of LPS-treated *Snca^+/+^* rats did not alter motor behavior, evaluated by rotarod and open field tests. This result may not be surprising because a high degree of neuronal degeneration is required for motor symptoms to be evident, in animals ^52^ and humans ^59,60^. Similarly, the decreased DA release in LPS *Snca^+/+^* rats did not alter the preference for a sweet-tasty solution, commonly used to assess the presence of a depressive-like behavior, such as anhedonia. In a recently published work, reduced sucrose consumption has been reported shortly after systemic LPS administration ^61^. However, this condition could reflect a more acute phase in response to LPS-induced inflammatory activation.

Our data support a dual-hit hypothesis for PD, whereby elevated endotoxin under a genetic predisposition may interact or synergize, driving the neurodegenerative processes that underlie the neuropathology of PD. Although we cannot discern whether the synergy between α-synuclein overexpression and LPS-induced inflammation is limited to an acceleration of PD-like symptoms typically expressed at later stages in *Snca*^+/+^ rats or exacerbates their expression, experiments performed at later stages of LPS injection are needed to clarify this problem. Another limitation of this study is to truly discriminate between infiltrated and parenchymal T cells and those present in the remaining blood vessels upon tissue dissociation, requiring a deeper histological investigation to overcome the difficulty to reliably determine the absolute abundance of T cells across the tissue.

In conclusion, our double-hit model could be relevant to familiar forms of PD, in which the development of a PD profile may depend on subjective exposure to risk environmental factors through inflammatory responses. More importantly, our findings provide additional evidence for the role of the immune system in playing a key role in the onset and progression of PD.

## MATERIAL AND METHODS

### Animals

Homozygous transgenic rats (Sprague-Dawley background) that overexpress the full-length human SNCA locus under the control of the endogenous human regulatory elements (*Snca^+/+^*) were used ^18^. Two or three rats per cage were housed under standard conditions (23 ± 1°C room temperature, 45-60% relative humidity, 12-h light/dark cycle) with food and water ad libitum. Following species-specific behavior, red plastic tubes and paper are added to the cages. All procedures follow the guidelines for the ethical use of animals from the Council Directive of the European Communities (2010/63/EU) and were approved by the Italian Ministry of Health (Authorization N°617-2019PR). Experimental animals were obtained by crossing heterozygous males with heterozygous female rats and were confirmed as WT or *Snca*^+/+^ following genotyping with quantitative PCR using DNA from ear biopsies and the primers for copy numbers of the α-synuclein transgene: SynProm-F: 5′-cgctcgagcggtaggaccgcttgttttagac-3′ and LC-SynPromR: 5′-cctctttc cacgccactatc-3′, normalized to the rat β-actin reference gene with primers: β-actin-F: 5′-agccatgtacgtagccatcca-3′ and β-actin-R: 5′-tctccggagtccatcacaatg-3′ ^18,19,21^.

### Treatment protocol

A single dose of lipopolysaccharide (LPS) from *Escherichia coli* O111:B4 (Sigma-Aldrich, #L2630, no less than 500,000 endotoxin units/mg) at the concentration of 5mg/kg dissolved in sterile saline (NaCl 0.9%) was intraperitoneally (i.p.) injected in *wild-type* (WT) (n=14) and *Snca^+/+^* (n=15) rats at two months of age ^62^. Control groups consist of a single i.p. injection of sterile saline in WT (n=15) and *Snca^+/+^* (n=16) rats at the same age. Animals were randomly divided into four experimental groups.

### Sickness Behavior and body weight

I.p. administration of LPS induces a sickness behavior in mice or rats ^30^. In this study, the signs consisting of absent exploration and locomotion, curled body posture, irregular fur, piloerection, and closed eyes were evaluated in the LPS-treated and control (saline) group over time after intraperitoneal injection of LPS, as previously described by ^30,31^. The animals were individually placed in a cage and scored on a four-point scale: 0 = no signs 1 = one sign, 2 = two signs, and 3 = three or more signs. The experimenter quantifying signs of sickness was blind to experimental and control conditions. Sickness behavior and body weight were monitored after 2 h, 18h, 24h, 42h, 48h, 72h, 96h, and 168h after the i.p. injection.

### Tissue collection

Immunohistochemical and cytofluorimetric analyses were carried out on brain tissues. To prevent blood cell contamination from SNpc and striatal tissues, brains were perfused and meninges were carefully removed prior to collection and analysis. Perfusion was carried out transcaridally with 1% heparin in 0.1 M sodium phosphate buffer (PB), after deep anesthesia (Rompum; 20mg/ml, 0.5 ml/kg; Bayer, and Zoletil; 100mg/ml, 0.5 ml/kg; Virbac). The brain was taken from the skull of the animal and the two hemispheres were divided into two halves along the medial sagittal line. The left hemisphere was treated for subsequent immunohistological analysis, and the right hemisphere was freshly dissected and processed for high dimensional flow cytometry. The left hemisphere was postfixed in 4% paraformaldehyde (PFA) in phosphate buffer (PB; 0.1 M, pH 7.4) at 4 ° C, kept for 72 hours in 4% PFA and replaced every 24 hours with a fresh solution.The samples were rinsed three times in a PB and immersed in 30% sucrose and 10% glycerol solution at 4 ° C until sinking for cryoprotection. Subsequently, the brains were embedded in OCT, frozen in isopentane, cooled into liquid nitrogen, and stored at −80 ° C. Coronal sections (30 μm) were cut from the anterior part of the brain to the midbrain using a cryostat (Leica) at −20°C. Slices were stored at 4°C in 0.1 M PB containing 0.02% sodiumazide before being further processed for immunohistochemical staining.

### Dissociation of the substantia nigra and striatum

After sacrifice of the animal, the substantia nigra and striatum were collected and immersed in D-PBS with high glucose buffer at 4 ° C. According to the manufacturer’s protocol, the tissue was immediately dissociated to single-cell suspension by enzymatic degradation and myelin removal using the adult brain dissociation kit and GentleMACS dissociator (Miltenyi Biotec.). Isolated tissues were placed in C-tubes, and a mixture containing Enzyme Pand Buffer Z was added to the samples. A second mix containing enzyme A and Buffer Y is then added, with the C tubes tightly closed and attached upside down onto the sleeve of the gentleMACS Octo Dissociator with Heaters (Miltenyi Biotec.). The appropriate program 37C_ABDK_02 was run for 30 minutes, and the c-tube was detached and centrifuged to collect the sample, which was then resuspended and filtered to a 70 um MACS SmartStrainer with 10 ml of cold D-PBS with high glucose. After centrifugation at 300 g for 10 minutes, the sample was resuspended in debris removal solution, gently overlayed in cold D-PBS with high glucose and centrifuged at 3000 g for 10 minutes with full acceleration brake. The two top phases were aspirated and discarded, and the sample was resuspended in cold D-PBS with highglucose, centrifuged at 1000 g for 10 minutes, and then resuspended in the staining buffer ready for cell count and flow cytometry staining.

### Immunophenotyping from immune cells of substantia nigra and striatum by high-dimensional flowcytometry

Cellular phenotypes were evaluated using multiparametric flow cytometry panels containing markers to identify cell types and activation states. These markers allowed us to exclude all cells of no interest on physical parameters (side and forward scatter) and to gate on specific cells of interest. Despite the deep perfusion and the careful removal of meninges, flow cytometry staining cannot discriminate infiltrating immune cells from those few cells that might still be present within the tissue vessels. For immunophenotyping of substantia nigra and striatum, single cell suspension was stained on the cell surface in different panels with PerCP5.5-conjugated anti-CD45 (1:100, BioLegend Cat# 103132 (also 103131), RRID:AB_893340), APC-Vio770-conjugated anti-CD11b/c (1:100, Miltenyi Biotec Cat# 130-121-054, RRID:AB_2752215), APC-conjugated anti-CD161 (1:100, BioLegend Cat# 205606, RRID:AB_11142680) or CD86 (1:100, Miltenyi Biotec Cat# 130-109-130, RRID:AB_2659403), PE-conjugated anti-CD45RA (1:100, BioLegend Cat# 202307, RRID:AB_314010), PE-Vio770-conjugated anti-CD3 (1:100, Miltenyi Biotec Cat# 130-103-773, RRID:AB_2657103) or anti-MHC-II (1:100, Miltenyi Biotec Cat# 130-107-822, RRID:AB_2652895), andFITC-conjugated anti-granulocytes (1:100 Miltenyi Biotec Cat# 130-108-084, RRID:AB_2651886) or VioBright-conjugated anti-ACSA-2/O4/CD31 (1:100, Miltenyi Biotec) for 15 minutes at 4°C in the dark. After staining, cells were washed and resuspended in PBS, ready to be acquired. These markers allowed us to correctly identify the different cell populations. Briefly, cells were first gated as CD45+ to identify the total leucocytes/resident immune cells and inside this gate, microglial cells were identified as CD45^low^CD11b/c^+^ cells and myeloid cells as CD45^high^CD11b^high^, which were subsequently identified as macrophages according to the expression of F4/80. The remaining of CD45^high^CD11b^low^ cells were further gated to identify T-lymphocytes as CD3^+^, B-lymphocytes as CD45RA^+^, and NK cells as CD161^+^. The expression of CD68 and MHC-II was further assessed within the microglial cell population. All samples were acquired on a 13-color Cytoflex (Beckman Coulter). For each analysis, at least 0.2×10^6^ live cells were acquired by gating on aqua Live/Dead negative cells and then analyzed using the Flowjo analysis software (FlowJo 10.0 (RRID:SCR_008520); Tre Star; https://www.flowjo.com/solutions/flowjo), as reported ^63^.

### Immunophenotyping of peripheral blood immune cells by flow cytometry

Peripheral blood was collected from the animals and whole blood was lyzed with the RBC (1X) solution for 10 minutes at 37°C. After lysis, cells were washed and resuspended in cold PBS for immunophenotyping. Cells were stained at the cell surface with FITC-conjugated CD4 (1:100, BioLegend Cat# 203406 (also 203405), RRID:AB_1227553) or anti-agranulocytes (1:100, Miltenyi Biotec Cat# 130-108-084, RRID:AB_2651886), PerCP5.5-conjugated anti-CD8 (1:100, BioLegend Cat# 201712, RRID:AB_2075263) or anti-CD45 (1:100, BioLegend Cat# 103132 (also 103131), RRID:AB_893340), APC-conjugated anti-CD25(1:80, BioLegend Cat# 202114 (also 202113), RRID:AB_2814102), PE-conjugated anti-CD45RA (1:100, BioLegend Cat# 202307, RRID:AB_314010), APC-conjugated anti-CD161 (1:100, Biolegend Cat# 205606, RRID:AB_11142680), PE-Vio770-conjugated anti-CD3 (1:100, Miltenyi Biotec Cat# 130-103-773, RRID:AB_2657103) and APC-Vio770-conjugated anti-CD11b/c (1:100, Miltenyi Biotec Cat# 130-121-054, RRID:AB_2752215) for15 minutes at 4°C in the dark. After staining, cells were washed and incubated with a fixing solution (1:3, BD Biosciences Cat# 554722, RRID:AB_2869010) in the dark for 15 minutes at room temperature. The cells were then washed with PBS and permeable with Cytofix/Cytoperm (1:10, BD Biosciences Cat# 554722, RRID:AB_2869010) and then stained intracellularly with APC-conjugated anti-IFN-γ (1:100, Biolegend Cat# 507810, RRID:AB_756180), PE-conjugated anti-IL-17 (1:30, eBioscience Cat# 12-7177-81, RRID:AB_763582) and PE-CF594-conjugated anti-FoxP3 (1:50, Biolegend Cat# 320008, RRID:AB_492980) in Cytoperm at room temperature for 30 min. After incubation, cells were washed and resuspended in PBS, ready to be acquired. These markers allowed us to correctly identify the different cell populations. Briefly, cells were then gated on CD45 to identify total leucocytes.Inside this gate, granulocytes were identified with their specific marker, monocytes and myeloid cells as CD11b/c^+^, T-lymphocytes as CD3^+^, B-lymphocytes as CD45RA^+^ and NK cells as CD161^+^ cells. All samples were acquired in a 13-color Cytoflex (Beckman Coulter) and for each analysis, at least 0.5×10^6^ live cells were acquired by gating on aqua Live/Dead negative cells and then analyzed by Flowjo analysis software (FlowJo 10.0 (RRID:SCR_008520); Tre Star; https://www.flowjo.com/solutions/flowjo), as reported ^19,63,64^.

### Immunohistochemistry

For stereological cell count and optical density analysis, SNpc and striatum slices were processed for chromogenic immunohistochemistry. First, endogenous peroxidase was neutralized with a 0.3% H_2_O_2_ solution in PB. The sections were then incubated with a blocking solution made with 5% normal goat serum (NGS) and 0.3% TritonX-100 (Sigma-Aldrich) in a PB solution for 1 h to prevent nonspecific binding of antibodies. Free-floating sections were incubated with the primary anti-TH antibody (1:1000, Millipore Cat# MAB318, RRID:AB_2201528) or anti-alpha synuclein (pS129) (1:500, Abcam Cat# ab51253, RRID: AB_869973) diluted in PB containing 0.3% Triton X-100 overnight at 4°C. After three rinses, sections were incubated with a biotinylated secondary antibody (Millipore Cat# AP124B, RRID:AB_92458) followed by incubation with extravidin-peroxidase (1:1000, Sigma-Aldrich Cat# e2886, RRID:AB_2620165). Antigen-antibody binding is then detected using chromogen 3,30-diaminobenzidine (DAB; Sigma-Aldrich). Finally, sections were dehydrated in a series of alcohols (50%, 70%, 95%, 100%), cleared in Xylene, and cover slipped with Entellan mounting medium (Sigma-Aldrich).

### Immunofluorescence

The sections were first incubated with a blocking solution made with 5% normal donkey serum and 0.3% TritonX-100 (Sigma-Aldrich) in a PB solution for 1 h to prevent nonspecific binding of antibodies. Primary antibodies (TH/DDC and NeuroTrace 435/455 (1:200, Thermo Fisher Scientific, Cat# N21479); TH/IBA1; TH/CD3; listed in Table 1) were incubated during the week in PB containing 0.3% Triton X-100 at 4 ° C and then incubated for 2h at room temperature with adequate secondary antibodies (listed in Table 1). The sections were counterstained with DAPI (4’,6-diamidin-2-fenilindolo). The specificity of immunofluorescence labeling was confirmed by omitting primary antibodies and instead using normal serum (negative controls). Confocal laser scanner microscopy (LSM800, Zeiss, Oberkochen, Germany) was used to acquire the images.

**Table 1.** Primary and secondary antibodies used for immunofluorescence.

| **Primary antibodies** | **Cat. Number** | **RRID** | **Host** | **Dilution** | **Manufacturer** |
| --- | --- | --- | --- | --- | --- |
| Tyrosine hydroxylase | #MAB318 | AB_2201528 | Mouse | 1:1000 | Millipore |
| IBA1 | #019-19741 | AB_839504 | Rabbit | 1:800 | Wako |
| DOPA Decarboxylase | #GTX134053 | AB_2887192 | Rabbit | 1:800 | Gene Tex |
| CD3 | #ab16669 | AB_443425 | Rabbit | 1:200 | Abcam |
| **Secondary antibodies** | **Cat. Number** |  | **IgG** | **Dilution** |  |
| Alexa Fluor 488 donkey | #A21202 | AB_141607 | Mouse | 1:200 | Thermo Fisher Scientific |
| Alexa Fluor 555 donkey | #A31572 | AB_162543 | Rabbit | 1:200 | Thermo Fisher Scientific |

### Stereological cell count

The stereological probe of the three-dimensional optical fractionator stereological probe was used to estimate the numberof TH^+^ and TH/DDC neurons in the SNpc and the Ionized calcium-binding adapter molecule 1-positive (IBA1^+^) cells in the SNpc and striatum. We used the stereo investigator system (MicroBright-Field, Vermont, USA). An optical microscope (Axio Imager M2, Zeiss, Oberkochen, Germany) equipped with a motorized stage, a MAC 6000 controller (Ludl Electronic Products, Ltd) and a camera are connected to software Stereoinvestigator 2019.1.3 (RRID:SCR_018948; https://www.mbfbioscience.com/stereo-investigator). The region of interest was defined by TH staining and the area distinction was performed according to the Paxinos and Franklin atlas of the rat brain and Franklin’s atlas and outlined with a 5-fold objective. The three-dimensional optical fractionator (x, y, z dimension of 80×80×30 μm, with a guard zone of 5 μm along the z axis) was used. The cells were marked with a 40x objective. Nine sections per animal, with five (SNpc) or six (dorsal striatum) sections intervals to cover the entire area, were counted. All quantifications were performed by an investigator blind to the experimental groups.

### Morphological analyses of tyrosine hydroxylase fibers

Three free-floating sections (30 μm) per animal, including the dorsolateral striatum and midbrain, were immunoistochemically stained for TH or pSyn (129). The sections were photographed with a light microscope (Axio Imager M2, Zeiss, Oberkochen, Germany). Densitometric analysis (OD) of TH^+^ fibers or pSyn (129) was analyzed using the Java image processing and plugin analysis program in ImageJ (NIH, USA; RRID:SCR_003070; https://imagej.net/). Densitometry values were corrected for nonspecific background staining by subtracting densitometric values from the cortex for the striatum and from the surrounding neuropil for midbrain areas in the same images.

TH optical density was measured in the dorsolateral striatum and SNpr, while immunoreaction levels of pSyn (S129) were evaluated in SNpc, VTA, pontine nuclei, and dorsolateral striatum.

All quantifications were performed by an investigator blind to the experimental groups.

### Sholl Analysis

IBA1^+^ cells were imaged with an optical microscope (Axio Imager M2, Zeiss, Oberkochen, Germany) equipped with a motorized stage and a camera connected to Neurolucida 2020.1.2 (MicroBright-Field, Vermont, USA; RRID:SCR_001775; http://www.mbfbioscience.com/neurolucida) that allows quantitative 3D analysis of the entire cell body and branching. Only cells with intact processes that were not obscured by background labeling or other cells were included in cell reconstruction. The area and perimeter, number of intersections, the number of nodes(branch points) and the endings, and total length of the processes were traced and exported to Neurolucida Explorer 2019.2.1 (MicroBright-Field, Vermont, USA; RRID:SCR_017348; https://www.mbfbioscience.com/neurolucida-explorer). To account for changes in cell complexity concerning distance from the soma, concentric rings (radii) were spaced 10 µm apart around the cell, centered on the centroid of thecell body and radiating outward.

Branching originating from the soma and the number of branch points and endings, processes intersecting the radii, and process length were measured as a function of the distance from the cell soma for each radius. In general, fifteen cells from five different sections of SNpc or striatum per animal were randomly selected for analysis, and all data were subsequently averaged for each rat ^19^. All quantifications were performed by an investigator blind to the experimental groups.

### Cortico striatal slice preparation

Acute brain slices were obtained after isoflurane anesthesia and decapitation. The brain was quickly removed, and 250–300 µm thick coronal slices containing the striatum were cut with a Leica vibratome (VT1200S) using chilled bubbled (95% O2, 5% CO2) ‘sucrose-based’ artificial CSF (aCSF) solution containing (in mM): KCl3, NaH_2_PO_4_ (1.25), NaHCO_3_ (26), MgSO_4_ (10), CaCl_2_ (0.5), glucose (25), sucrose (185); ∼300 mOsm, pH 7.4). Slices were used after a recovery period of at least 40 min in standard aCSF solution containing (in mM): NaCl 126, KCl 2.5, NaH2PO_4_ (1.2), NaHCO_3_ (24), MgCl_2_ (1.3), CaCl_2_ (2.4), glucose (10) (∼290 mOsm, pH 7.4) at 32 °C.

### Constant Potential Amperometry

Amperometric detection of electrically evoked DA release was performed in acute brain slices that contained the striatum. The carbon fiber electrode was gently placed in the tissue, and the voltage (MicroC potentiostat, World Precision Instruments) was imposed between the carbon fiber electrode and the Ag/AgCl pellet at 0.60 V. The DA-recording carbon fiber electrode (diameter 30 μm, length 100 μm, World Precision Instruments) was positioned near a bipolar Ni/Cr stimulating electrode to a depth of 50–150 μm into the coronal slice. For stimulation, a single rectangular electrical pulse was applied using a DS3 stimulator (Digitimer) every 5 min along a range of stimulation intensities (20–1000 μA, 20–80 μs duration). Under stimulation, DA-containing vesicles were released from the presynaptic terminals. When DA molecules in the synaptic milieu hit the carbon surface, electrodes were transferred and a current can be measured. In response to a protocol of increasing stimulation, a plateau of DA release was reached at maximum stimulation (1000 μA,80 μs). To measure changes in DA synaptic activity caused by drugs that affect dopamine transporter (DAT) or D23 receptors, we analyzed the effects on evoked DA release by applying cocaine or quinpirole, respectively. Drugs were superfused to the striatal slice when the extracellular DA response to electrical stimulation was stable for four or five successive stimulations. The evoked release of DA was evaluated after bath perfusion of 0.3 o1 μM Cocaine dissolved in aCSF ^35^. The perfusion was carried out for 10 minutes when the maximum effect on DAT by cocaine was reached, and then it was washed and followed for at least 1 h. The activity of D2 receptors on DA release was evaluated after bath perfusion of 0.03 μM quinpirole, an agonist of the D2 receptor. For cocaine, quinpirole was perfused for 5 minutes and washed for 30 minutes. The signals were digitized with Digidata 1440A coupled to a computer running pClamp 10 (Molecular Devices; RRID:SCR_011323; http://www.moleculardevices.com/products/software/pclamp.html). At the end of each experiment, electrode calibration was performed with bath-perfused DA at known concentrations (0.3–10 μM).

### Behavioral tests

Three months after single injection, rats were tested for Rotarod, open field, and sucrose preference tests.

The animals were accustomed for 1 h to the testing room before the beginning of the Rotarod and open field tests. At the end of each session/trial, all the apparatus was cleaned with 70% ethanol to remove olfactory signals.

### Rotarod test

Motor coordination and balance were tested using an accelerating Rotarod protocol (LE8355, Panlab, Spain). The apparatus consisted of a rod suspended horizontally at 47 cm from the floor. All animals were accustomed to being placed on the rod rotating at low speed (4 rpm) for at least 30 seconds before the first session of the test, which consisted of four sessions. Sessions were performed on two consecutive days (with an intersession interval of 3.5 h) with an accelerating rod from 4 to 40 rpm in 300 s. Each session consisted of three trials, with at least 5 min of rest between the trials. For each trial, the time the rat fell from the rod was recorded (maximum 300 s). The first three sessions are considered pre-training on the Rotarod apparatus to reach a stable performance. The fourth session was considered the final test and the latency to fall from the rod was analyzed by averaging the three trial records.

### Open field test

The open field is a simple test to evaluate general motor and exploratory behavior in rodent models of CNS disorders. The apparatus consisted of a cylindrical plastic arena (100 cm in diameter) with dark walls (35 cm high) and floor, placed in a room with dim lighting. During the test, each animal was placed in the center of the arena and allowed to explore the apparatus for 10 min. Experiments were recorded with a video camera suspended above the arena and data were analyzed with EthoVision XT15 (Noldus, The Netherlands; RRID:SCR_000441; https://www.noldus.com/ethovision) video tracking software and the total distance traveled was measured.

### Sucrose preference test

The sucrose preference test assesses the animal’s interest in a sweet-tasting sucrose solution relative to unsweetened water. Rats were habituated to drinking from two equally accessible bottles and were singly housed for the test duration. The test cage was the same throughout the experiment. The rats were exposed to a double choice drinking test, with one bottle containing one sucrose solution (1%) and the other drinking water. To avoid side preference, the sucrose bottle was placed on the left for half of the cages and on the right for the other cages. Each bottle was refilled with fresh water or sucrose solution and re-inverted to its original position every day for two consecutive days. After 72h, the consumption of water and sucrose solution was measured with a graduated cylinder. A 48-hour testing period allowed us to exclude any effects of neophobia, artifact bias toward any one side, and perseveration effects. In addition, it provided information on long-term access to a rewarding stimulus. The percentage of sucrose intake and water intake was evaluated as follows:

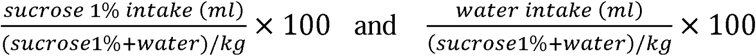

### Statistical analysis

All statistical analysis was performed with GraphPad Prism 8.0.1 (RRID:SCR_002798; http://www.graphpad.com/) using the appropriate statistical tests, as indicated in the legend of each figure. Data were normalized using the Shapiro-Wilk test or the Kolmogorov-Smirnov test, followed by the appropriate statistical test. The values of P <0.05 were considered to be statistically significant. All data were presented as mean ± SEM, and each point represents individual experiments.

## DATA AVAILABILITY STATEMENT

More information and request for resources and reagents should be directed to and will be fulfilled by the corresponding authors, Mariangela Massaro Cenere, and Nicola Biagio Mercuri

The complete data sets can be found in the Zenodo repository: 10.5281/zenodo.13969544

A detailed description of the conducted protocol is available at the protocols.io repository: (DOI: dx.doi.org/10.17504/protocols.io.j8nlkoj65v5r/v1 (Private link for reviewers: https://www.protocols.io/private/EB5CE11EBE8F11EE8E330A58A9FEAC02 to be removed before publication)

## AKNOWLEDGEMENTS

This research was funded in whole or in part by Aligning Science in Parkinson’s [020505] through the Michael J. Fox Foundation for Parkinson’s Research (MJFF). For the purpose of open access, the author has applied a CC BY public copyright license to all Author Accepted Manuscripts arising from this submission. Conducted work was also supported by NEXTGENERATIONEU (NGEU) and funded by the Ministry of University and Research (MUR), National Recovery and Resilience Plan (NRRP), project MNESYS (PE0000006) – A Multiscale integrated approach to the study of the nervous system in health and disease (DN. 1553 11.10.2022), and by the Ministry of University and Research (MIUR) - PRIN (Bando 2017, Prot. 2017ENN4FY, N.B.M). The sponsor did not play a role in study design, data collection, analysis, and interpretation of data, or writing of this manuscript.

## AUTHOR CONTRIBUTIONS

MMC, VC, and NBM designed the experiments, MMC performed i.p. injections and sacrifices. N.C. provided transgenic animals. MMC, MT, SLD, MF, CG, AM, FC, and BZ collected data and performed data analysis with feedback from EP, DC, AL, LP, NB, FRF, and VC. MMC, VC, and NBM wrote and revised the manuscript. All authors contributed to the article and approved the submitted version.

