## Supplementary material for "Systemic inflammation accelerates neurodegeneration in a rat model of Parkinson’s disease overexpressing human alpha synuclein": Supplementary data.docx

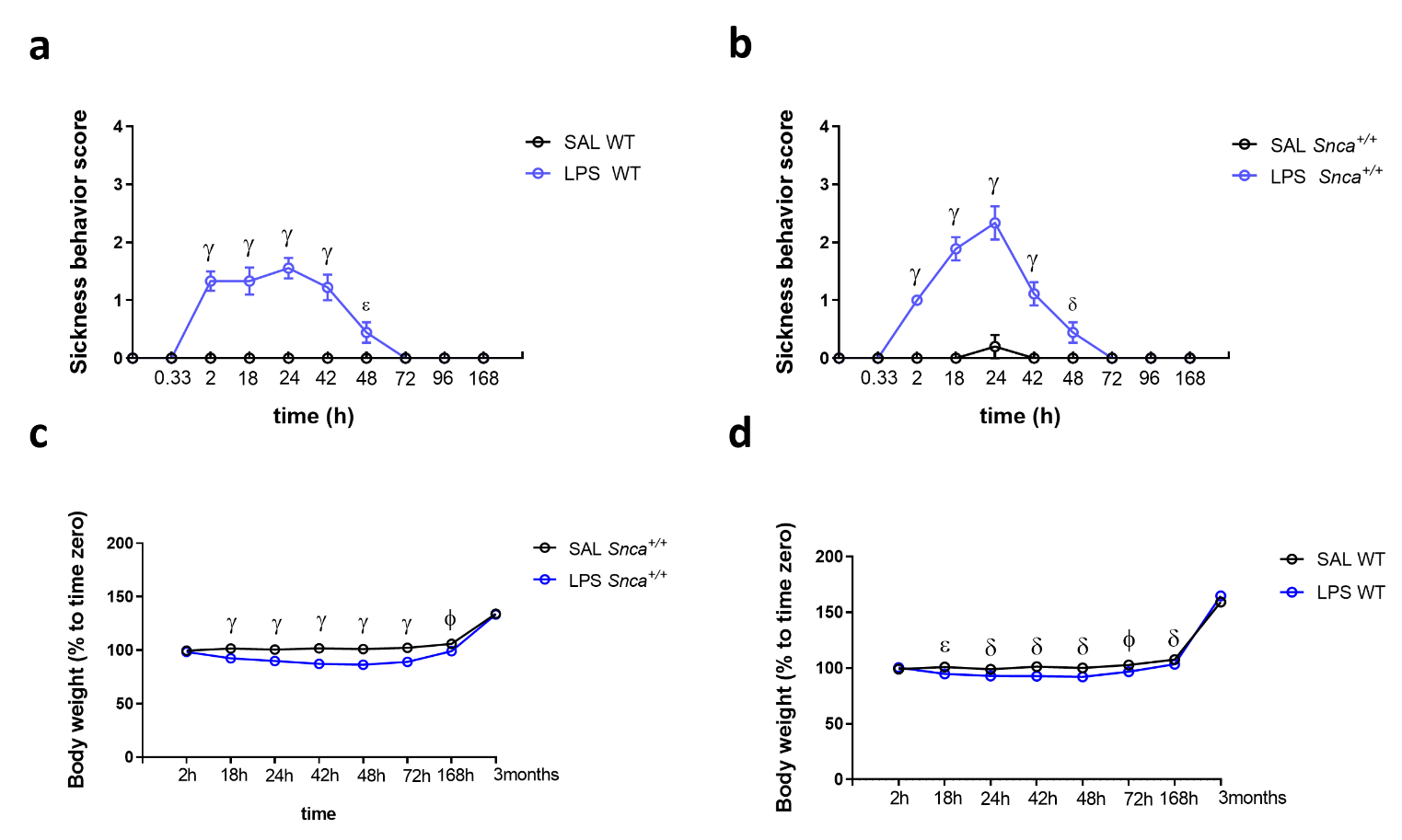

**Supplementary Figure 1. Time course of sickness behavior and body weight. a-b)** Clinical evolution of sickness behavior of WT (A) and *Snca^+/+^* (B) rats after a single intraperitoneal (i.p.) injection of LPS or saline. All values represent the mean ± SEM. (n=10 SAL WT, n=9 LPS WT, n=10 SAL *Snca^+/+^*, n=9 LPS *Snca^+/+^*; **a:** Two-way repeated measure ANOVA Time vs Treatment; Interaction, F (9, 153) = 30.99, p<0.0001, Time, F (9, 153) = 30.99, p<0.0001, Treatment, F (1, 17) = 90.41, p<0.0001; ^ε^p<0.01, ^γ^p<0.0001, with Bonferroni’s *post hoc* multiple comparison test; **b:** Two-way repeated measure ANOVA Time vs Treatment; Interaction, F (9, 153) = 36.63, p<0.0001, Time, F (9, 153) = 44.42, p<0.0001, Treatment, F (1, 17) = 88.98, p<0.0001; ^δ^p<0.01, ^γ^p<0.0001, with Bonferroni’s *post hoc* multiple comparison test). **c-d)** Body weight changes following LPS i.p. administration in WT (C) and *Snca^+/+^* (D) rats All values represent the mean ± SEM. (n=10 SAL WT, n=9 LPS WT, n=10 SAL *Snca^+/+^*, n=9 LPS *Snca^+/+^*; **c:** Two-way repeated measure ANOVA Time vs Treatment; Interaction, F (7, 119) = 11.82, p<0.0001, Time, F (1.865, 31.70) = 1024, p<0.0001, Treatment, F (1, 17) = 8.885, p=0,0084; ^δ^p<0.01, ^ε^p<0.01, ^ϕ^p<0.001, with Bonferroni’s *post hoc* multiple comparison test; **d:** Two-way repeated measure ANOVA Time vs Treatment; Interaction, F (7, 119) = 43,90, p<0.0001, Time, F (2.024, 34.42) = 1001, p<0.0001, Treatment, F (1, 17) = 185.4, p<0.0001; ^ϕ^p<0.001, ^γ^p<0.0001, with Bonferroni’s *post hoc* multiple comparison test).

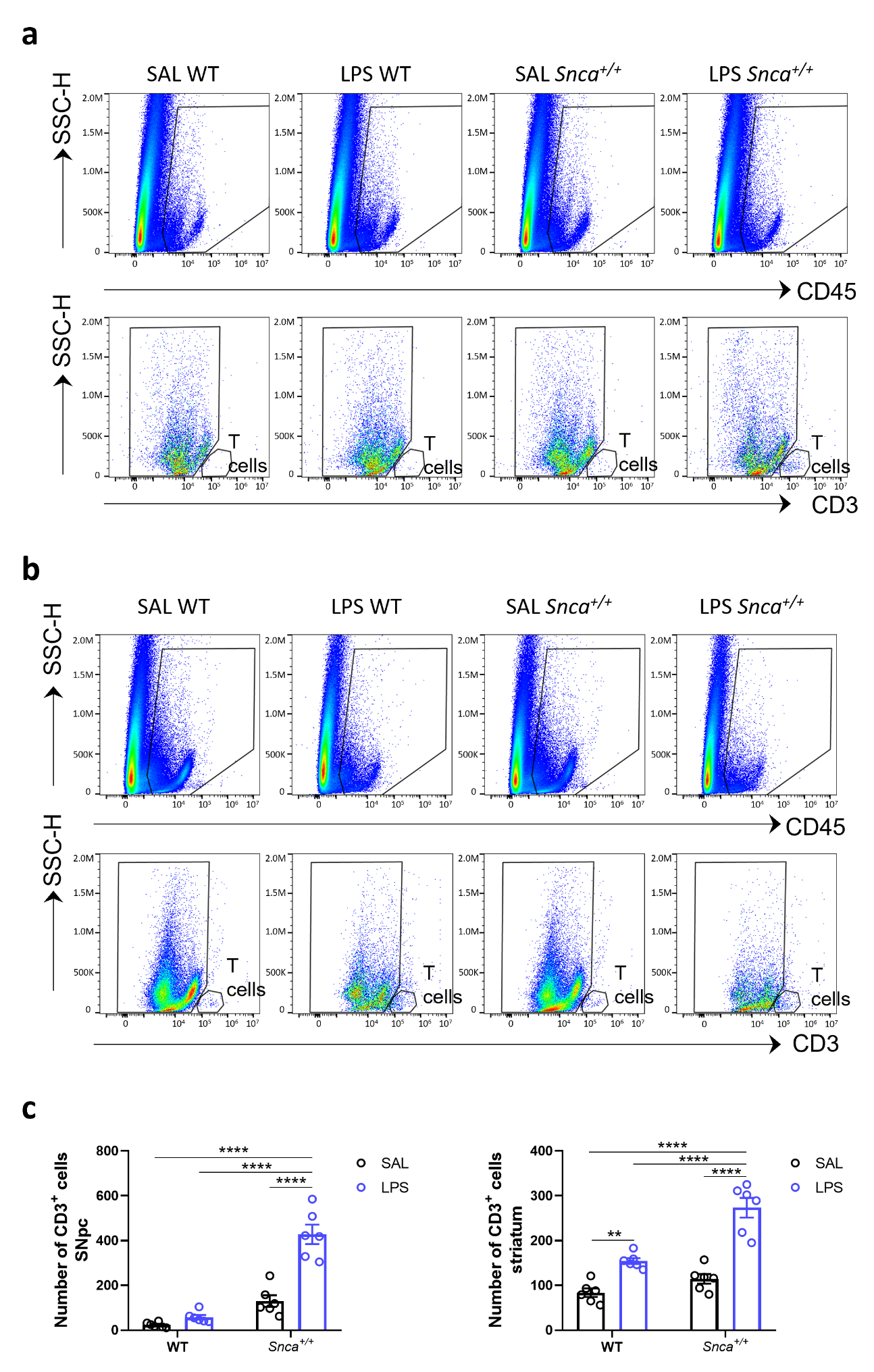

**Supplementary Figure 2. Gating strategy and T cell count. a-b)** Representative flow cytometry dot plots of isolated cells illustrating the gating strategy to identify total leukocytes based on CD45 and T cells based on CD3 in the SNpc (**a**) and striatum (**b**). **c)** Cell count of total CD3^+^ T cells in the SNpc (left) and in the striatum (right). Error bars represent ±SEM. (n=6 SAL WT, n=6 LPS WT, n=6 SAL *Snca^+/+^*, n=6 LPS *Snca^+/+^*; Ordinary two-way ANOVA for Genotype vs Treatment; SNpc: Interaction, F (1, 20) = 26.03, p<0.0001; Genotype, F (1, 20) = 85.29, p<0.001; Treatment, F (1, 20) = 41.38, p<0.0001; ****p<0.0001, with Bonferroni’s *post hoc* multiple comparisons test; striatum: Interaction, F (1, 20) = 10.70, p=0.0038; Genotype, F (1, 20) = 30.89, p<0.0001; Treatment, F (1, 20) = 72.47, p<0.0001; **p<0.01, ****p<0.0001, with Bonferroni’s *post hoc* multiple comparisons test).

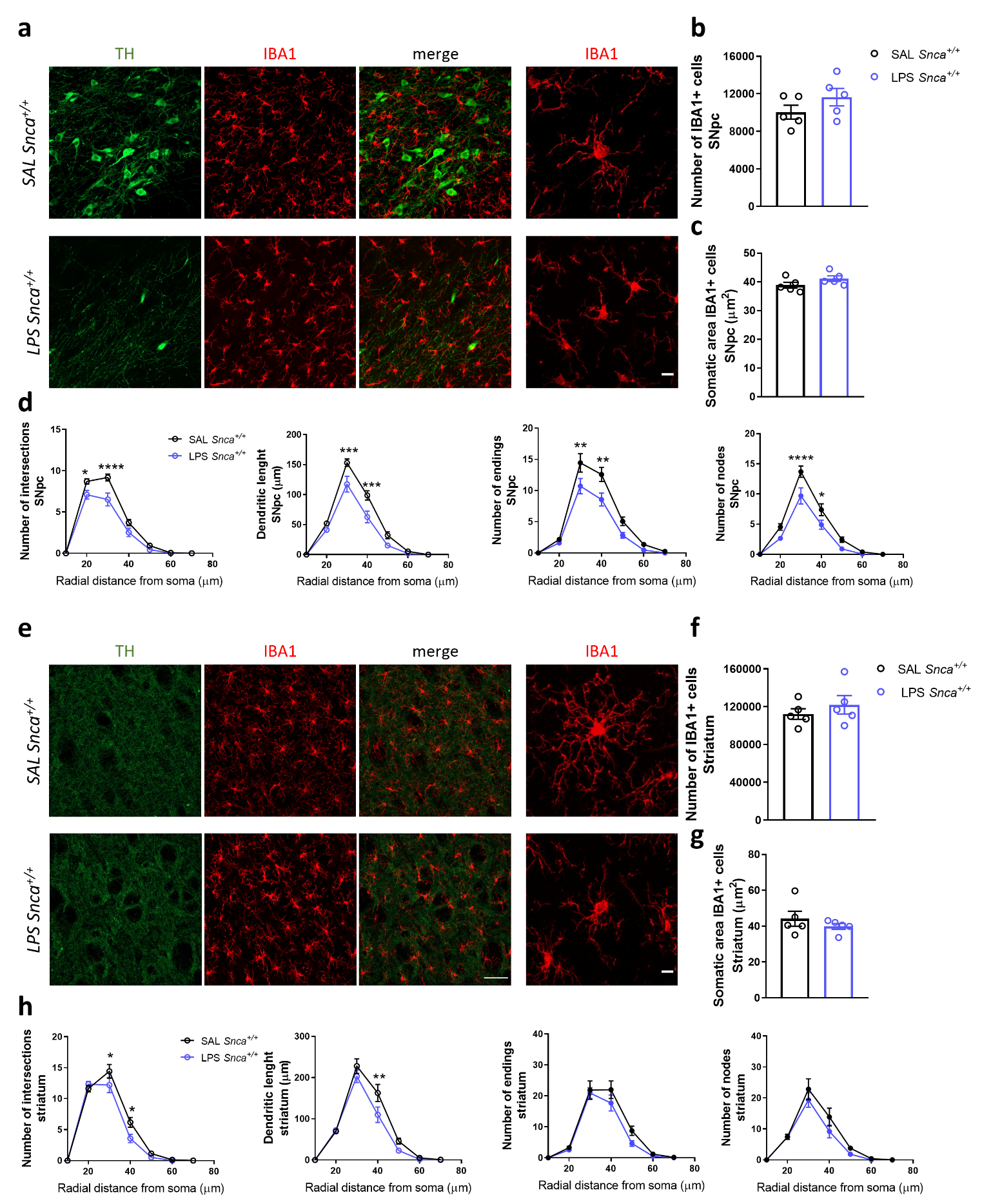

**Supplementary Figure 3. Microglial morphological changes reflecting its activation in CNS 3 months after LPS administration. a)** Left, representative confocal images of SNpc TH^+^ neurons (green) and IBA1^+^ microglial cell (red) and, right, relative micrograph showing an IBA1^+^ microglial cell (red) converted to a maximum intensity Z-stack projection (scale bar: 50µm, 10 µm). **b)** Bar plot of stereological IBA^+^ microglial cell unbiased count in the SNpc (n= 5 rats/group; Unpaired *t*-test, t=1.340, p=0.2172). **c)** Bar plot of IBA^+^ somatic area in the SNpc (n= 5 rats/group, Unpaired *t*-test, t=1.783, p=0,124). **d)** Panels indicating mean values of the number of intersections, distance length, number of ending and nodes at each radial distance from soma (μm), in the SNpc (n= 5 rats/group; Intersections: two-way ANOVA, Interaction, F (6, 56) = 4.408, p=0.0010; Treatment, F (1, 56) = 21.82, p<0.0001; distance, F (6, 56) = 225.2, p<0.0001; 20μm *p=0.0107, 30μm ****p<0.0001, with Sidak’s *post hoc* multiple comparison test; Length: two-way ANOVA, Interaction, F (6, 56) = 3.911, p=0.0025; Treatment, F (1, 56) = 24.22, p<0.0001; distance, F (6, 56) = 165.4, p<0.0001; ; 20μm ***p=0.0002, 30µm ***p=0.0002, with Sidak’s *post hoc* multiple comparison test; Endings: two-way ANOVA, Interaction, F (6, 56) = 2.749, p=0.0206; Treatment, F (1, 56) = 19.17, p<0.0001; distance, F (6, 56) = 105.7, p<0.0001; 30µm **p=0.0035, 40µm **p=0.0014, with Sidak’s *post hoc* multiple comparison test; Nodes: two-way ANOVA, Interaction, F (6, 56) = 3.203, p=0.0089; Treatment, F (1, 56) = 21.43, p<0.0001; distance, F (6, 56) = 108.1, p<0.0001; 30µm ****p<0.0001, 40µm *p=0.0292, with Sidak’s *post hoc* multiple comparison test). **e)** Left, representative confocal images of striatum TH^+^ neurons (green) and IBA1^+^ microglial cell (red) and, right, relative micrograph showing an IBA1^+^ microglial cell (red) converted to a maximum intensity Z-stack projection (scale bar: 50µm, 10 µm). **f)** Bar plot of stereological IBA^+^ microglial cell unbiased count in the striatum (n= 5 rats/group, Unpaired t-test, t=0.8688, p=0.4102). **g)** Bar plot of IBA^+^ somatic area in the striatum ( n= 5 rats/group, Unpaired t-test, t=0.9311, p=0.3790). **h)** Panels indicating mean values of the number of intersections, distance length, number of ending and nodes at each radial distance from soma (μm), in the striatum (n= 5 rats/group; Intersections: two-way ANOVA; Interaction, F (6, 56) = 2.452, p=0.0355; Treatment, F (1, 56) = 5.403, p=0.0238; distance, F (6, 56) = 219.9, p<0.0001; 30μm *p=0.0483, 40 µm *p=0.0119, with Sidak’s *post hoc* multiple comparison test; Length: two-way ANOVA; Interaction, F (6, 56) = 1.981; Treatment, F (1, 56) = 7.098, p=0.0101; distance, F (6, 56) = 131.5, p<0.0001; 40µm **p=0.0038, with Sidak’s *post hoc* multiple comparison test; Endings: two-way ANOVA; Interaction, F (6, 56) = 0.7716; Treatment, F (1, 56) = 3.907; distance, F (6, 56) = 79.87, p<0.0001; Nodes: two-way ANOVA, Interaction, F (6, 56) = 0.8197; Treatment, F (1, 56) = 3.468; distance, F (6, 56) = 55.65, p<0.0001). **b-d, f-h)** All data are mean values of the number of intersections, ending, nodes and distance length at each radius ± SEM.

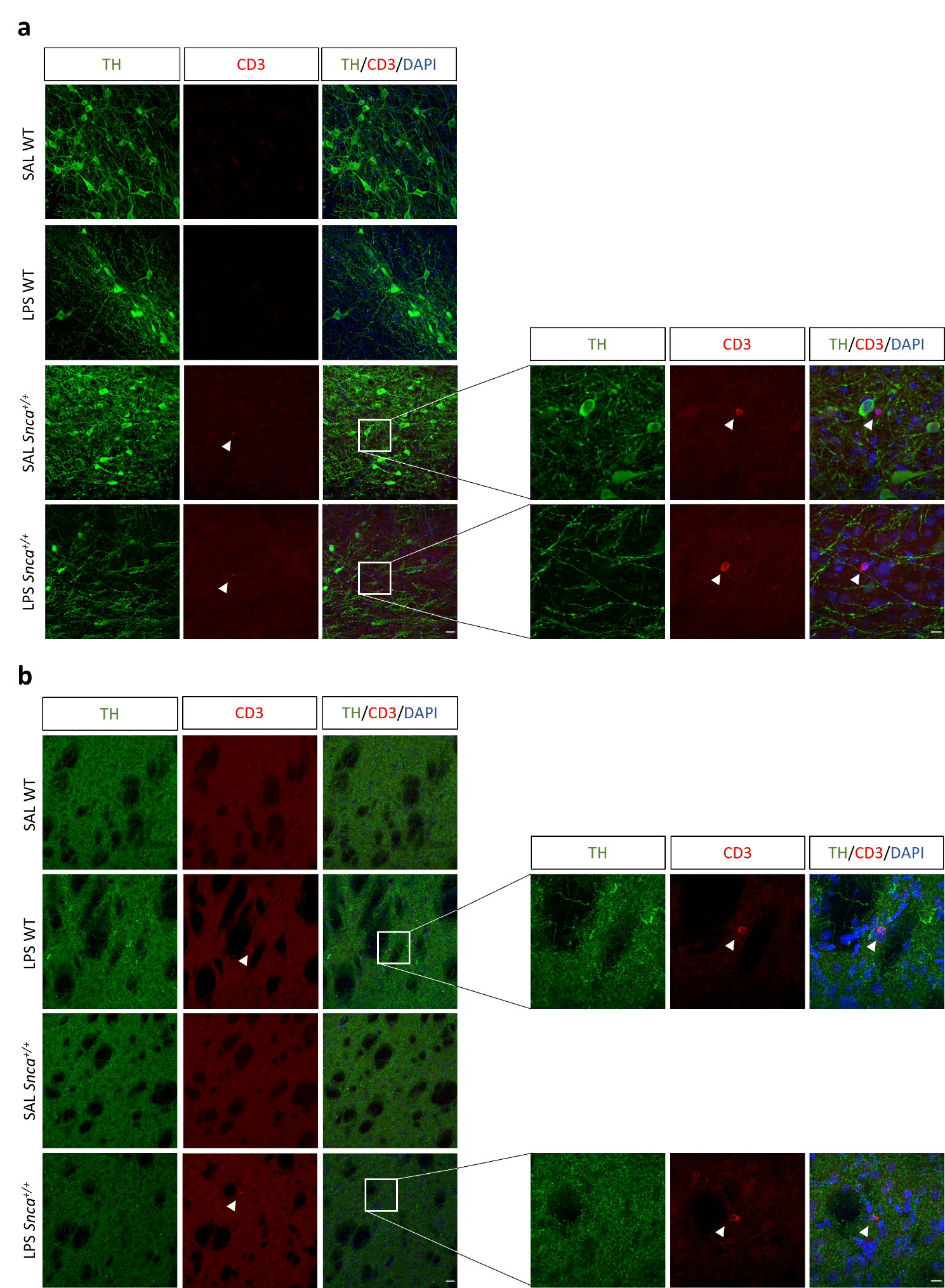

**Supplementary Figure 4. CD3^+^ cells in the SNpc and striatum. a)** Representative confocal images of CD3^+^ cells (red) and TH^+^ neurons (green) and relative micrographs of SNpc coronal sections (scale: 20μm; 10μm). The white arrowheads show T-lymphocytes nearby to TH^+^ neurons in the SNpc in SAL- and LPS-treated *Snca^+/+^* rats. Images are shown as maximum intensity Z-stack projection. **b)** Representative confocal images of CD3^+^ cells (red) and TH^+^ neurons (green) and relative micrographs of striatal coronal sections. White arrowheads show T-lymphocytes in LPS-treated WT and *Snca^+/+^* rats (scale: 20μm; 10μm). Images are shown as maximum intensity Z-stack projection.

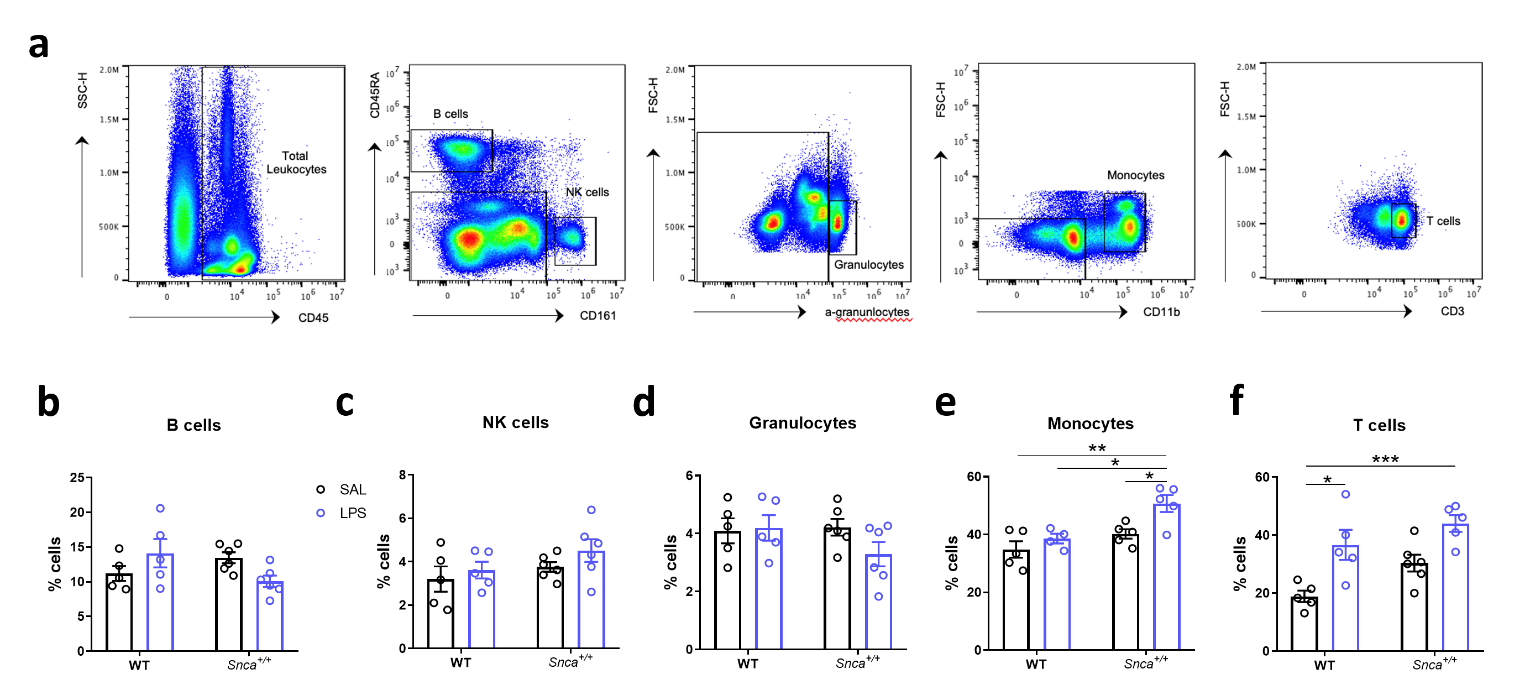

**Supplementary Figure 5. Peripheral blood mononuclear cell levels change 3 months after LPS administration.** **a)** Representative flow cytometry dot plots of isolated peripheral blood mononuclear cells illustrating the gating strategy to determine each cell population based on their expression levels of specific markers. **b-f)** Bar plots showing the percentages of B cells (**b**), NK cells (**c**), granulocytes (**d**), monocytes (**e**) and T-lymphocytes (**f**) in the peripheral blood of WT and *Snca^+/+^* rats (n=5 SAL WT, n=5 LPS WT, n=6 SAL *Snca^+/+^*, n=6 LPS *Snca^+/+^*, Ordinary two-way ANOVA for Genotype vs Treatment; **b**: Interaction, F (1, 18) = 6.680, p=0.0187, Genotype, F (1, 18) = 0.5326, Treatment, F (1, 18) = 0.03933; **c**: Interaction, F (1, 18) = 0.1479; **d**: Interaction, F (1, 18) = 1.707). **e**: Interaction, F (1, 15) = 1.921, Genotype, F (1, 15) = 12.80, p=0.0027; Treatment, F (1, 15) = 8.545, p=0.0105; *p<0.05, **p<0.01, with Bonferroni’s *post hoc* multiple comparisons test). **f:** Interaction, F (1, 17) = 0.3601; Genotype, F (1, 17) = 7.624, p=0.0134; Treatment, F (1, 17) = 21.07, p=0.0003; *p=0.0135, ***p=0.0005, with Bonferroni’s *post hoc* multiple comparisons test).

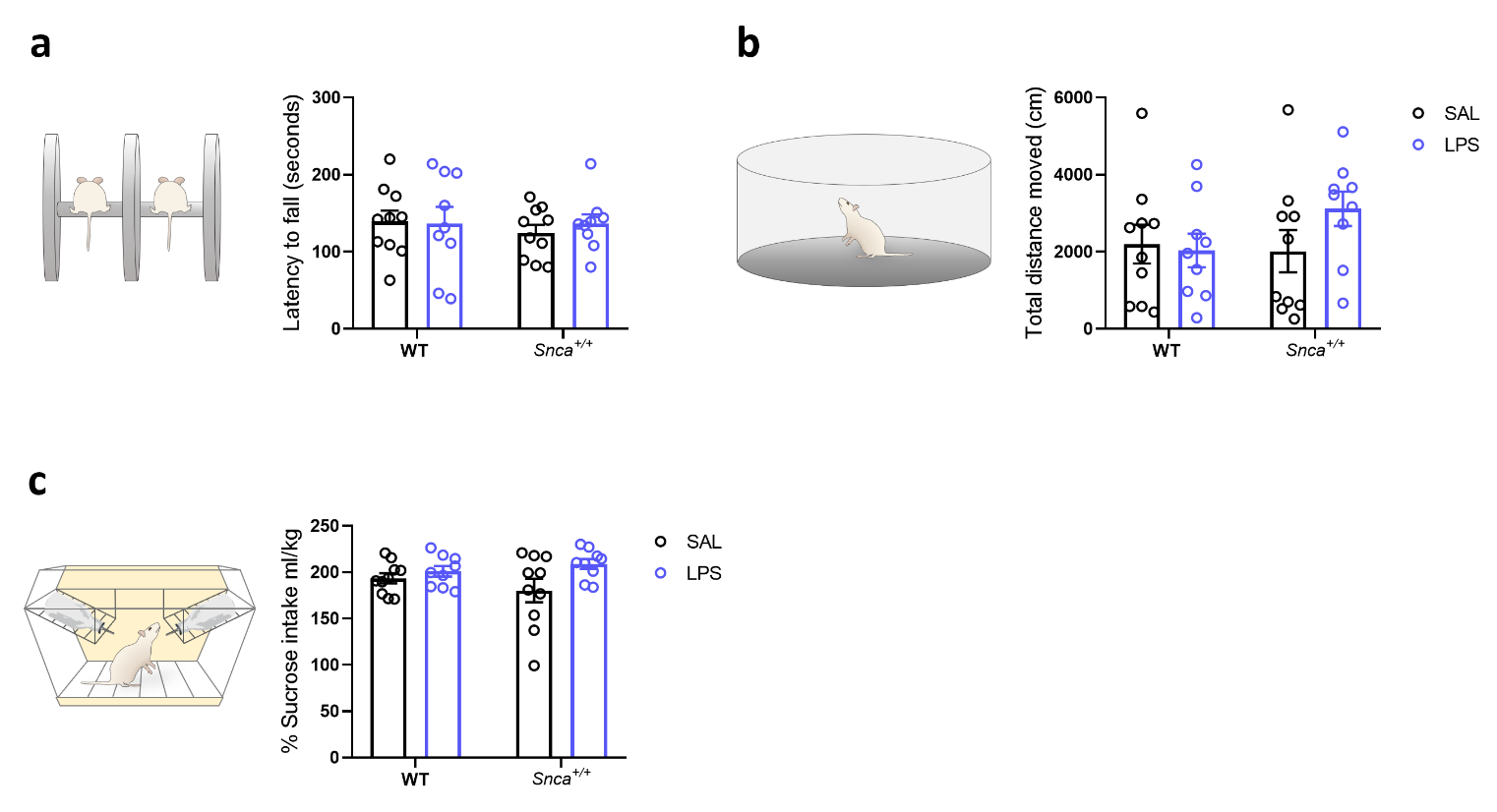

**Supplementary Figure 6. LPS does not alter general locomotor and depressive-like behavior 3 months after administration. a)** Left, schematic representation of the Rotarod test and its relative histogram showing latency to fall (in seconds). Each point represents mean values at the last trial ±SEM. (n=10 SAL WT, n=9 LPS WT, n=10 SAL *Snca^+/+^*, n=9 LPS *Snca^+/+^*; Ordinary two-way ANOVA for Genotype vs Treatment; Interaction, F (1, 34) = 0.2430). **b)** Left, schematic representation of open-field arena and its relative histogram showing total distance moved. Each point represents the actual value ±SEM. (n=10 WT SAL, n=9 WT LPS, n=10 *Snca^+/+^* SAL, n=9 *Snca^+/+^* LPS; Ordinary two-way ANOVA for Genotype vs Treatment; Interaction, F (1, 34) = 1.654). **c)** Left, schematic representation of sucrose preference test and its relative histogram showing the percentage of sweet solution consumption (n=10 SAL WT, n=9 LPS WT, n=10 SAL *Snca^+/+^*, n=9 LPS *Snca^+/+^*; Ordinary two-way ANOVA for Genotype vs Treatment; Interaction, F (1, 34) = 1.693).

|  | **Sample** | **Total Markers Counted** | **Number of Sections** | **Measured Defined Mounted Thickness** | **Estimated Population using Mean Section Thickness** | **Coefficient of Error (Gundersen), m=0** | **Coefficient of Error (Gundersen), m=1** |
| --- | --- | --- | --- | --- | --- | --- | --- |
| SAL *Snca+/+* | 1 | 188 | 9 | 28,5 | 9537,62 | 0,08 | 0,07 |
|  | 2 | 317 | 9 | 28,2 | 11784,47 | 0,06 | 0,06 |
|  | 3 | 250 | 9 | 29,2 | 8043,54 | 0,08 | 0,06 |
|  | 4 | 280 | 9 | 29,6 | 9140,81 | 0,08 | 0,06 |
|  | 5 | 227 | 9 | 29,0 | 11724,72 | 0,07 | 0,07 |
| LPS *Snca+/+* | 1 | 281 | 9 | 28,8 | 14402,14 | 0,06 | 0,06 |
|  | 2 | 205 | 9 | 27,9 | 10176,74 | 0,09 | 0,07 |
|  | 3 | 248 | 9 | 28,7 | 12658,19 | 0,09 | 0,06 |
|  | 4 | 190 | 9 | 28,9 | 9764,67 | 0,09 | 0,07 |
|  | 5 | 228 | 9 | 29,4 | 11914,55 | 0,07 | 0,07 |

**Supplementary table 1. Data Stereological IBA1^+^ cell count in SNpc**

|  | **Sample** | **Total Markers Counted** | **Number of Sections** | **Measured Defined Mounted Thickness** | **Estimated Population using Mean Section Thickness** | **Coefficient of Error (Gundersen), m=0** | **Coefficient of Error (Gundersen), m=1** |
| --- | --- | --- | --- | --- | --- | --- | --- |
| SAL Snca+/+ | 1 | 268 | 9 | 28,0 | 107288,09 | 0,08 | 0,06 |
|  | 2 | 333 | 9 | 27,5 | 130713,45 | 0,08 | 0,06 |
|  | 3 | 280 | 9 | 29,1 | 116304,05 | 0,08 | 0,06 |
|  | 4 | 282 | 9 | 29,1 | 117050,08 | 0,06 | 0,06 |
|  | 5 | 234 | 8 | 28,9 | 96548,37 | 0,09 | 0,07 |
| LPS Snca+/+ | 1 | 276 | 9 | 28,1 | 110827,99 | 0,07 | 0,06 |
|  | 2 | 298 | 9 | 27,5 | 116967,39 | 0,08 | 0,06 |
|  | 3 | 319 | 9 | 27,5 | 125323,02 | 0,08 | 0,06 |
|  | 4 | 258 | 8 | 27,2 | 100062,89 | 0,09 | 0,06 |
|  | 5 | 378 | 9 | 29,1 | 157191,14 | 0,07 | 0,05 |

**Supplementary table 2. Data from Stereological IBA1^+^ cell count in striatum**
